# Deciphering Mechanistic Signatures in Drug-Drug Interactions with Dual Topology Graphs

**DOI:** 10.64898/2026.09.23.753684

**Authors:** Wenjian Ma, Xiangpeng Bi, Huasen Jiang, Weigang Lu, Jie Nie, Shutan Lin, Jiaxin Lin, Zhiqiang Wei, Henggui Zhang, Shugang Zhang

## Abstract

Drug-drug interactions (DDIs) represent a critical challenge in drug development and clinical practice, as they can lead to severe adverse effects, including toxicity and reduced therapeutic efficacy. Deep learning methods have shown promise in large-scale, rapid DDI prediction; however, current approaches suffer from significant limitations in providing mechanistic insights into these interactions. Here, we propose DualTopoDDI, a dual-topology-enhanced interpretable deep learning model for DDI prediction. We applied DualTopoDDI to predict 9.2 billion potential interactions among approved drugs, achieving 97.99% high-confidence predictions. The model demonstrates molecular-level interpretability, identifying key substructures responsible for drug actions in terms of both atom-centric and bond-centric views. DualTopoDDI also excels in elucidating particular DDI toxicity mechanisms, which is validated by its successful explanation of the controversial cardiac toxicity in two COVID-19 drug combination regimens at the time. Evaluations across 11 benchmark datasets demonstrates that DualTopoDDI not only achieves state-of-the-art performance but also showcases robust generalizability and good interpretability. Overall, DualTopoDDI offers a powerful, interpretable tool for understanding and predicting drug-drug interactions, providing critical insights for drug safety and design.

## Introduction

The complexity of pathological mechanisms and the multi-morbidity caused by population aging clinically necessitate polypharmacy^1^. However, over 30% of adverse drug reactions (ADRs), such as cardiac arrhythmias and respiratory paralysis, are attributed to drug-drug interactions (DDIs)^2^, resulting in annual medical costs ranging from $5 to $7 billion^3,4^. Consequently, preemptive DDI risk assessment prior to polypharmacy is critical for safeguarding patient safety and conserving public health-care resources. On the other hand, the efficiency and costs of conventional pharmacological experimentation are far from satisfactory—even considering only whether ≈2,600 marketed drugs exhibit any toxicity (i.e., ignoring specific toxicity types), there are nearly 6.8 million drug-pair tests to perform^5,6^. Assuming a cost of $1,000 per pair and one day per test, evaluating all drug pairs would cost $6.8 billion and take up to 18,630 years. This crucial gap is also evidenced by the fact of frequently reported withdrawals of marketed drugs like terfenadine^7^. Therefore, an accurate and efficient computational measurement is needed for high-throughput evaluation of potential DDIs. In this context, Deep learning has emerged as a key tool to achieve this goal due to its ability to process high-dimensional molecular data and capture complex nonlinear interactions between drugs^8,9^.

A typical class of deep learning methods is to leverage biological knowledge graphs^10-12^. In this regard, multi-scale biomedical data is collected to construct knowledge graphs with drugs, targets, and diseases as entities. Variants of graph neural networks (GNNs)^13^ are then employed to infer unknown DDIs from known entity relationships. Although these methods achieve superior performance in DDI prediction tasks by incorporating external biomedical knowledge, they are less suitable for early-stage drug discovery scenarios where only molecular structural information is available. Meanwhile, substructure-based methods have been progressively proposed^14-17^. Such kind of methods are grounded in the chemical hypothesis that DDIs are mediated by key substructures or functional groups, thereby eliminating the need for external biomedical knowledge. By inputting a pair of drug molecules, the models can automatically extract substructural features to determine whether the drug pair interacts. Notably, multiple substructure extraction strategies are utilized in this process. For instance, drug molecules can be decomposed into non-redundant substructures (e.g., carboxylic acid groups, benzene rings, amine groups) using predefined chemical rules, thereby constructing a substructure library^14,15^. Subsequently, drug pairs are subjected to substructure matching against this library to generate global molecular representations. Another strategy involves using a message-passing framework to dynamically assign atomic importance weights on the molecular graph, indirectly identifying both known and unknown substructures^16-18^. Regardless of the strategy utilized, these substructure-based methods exhibit superior predictive performance on DDI tasks by capturing critical substructure information of drugs.

In brief, Deep learning technologies has emerged as a high-throughput and accurate alternative for DDI prediction. However, there are still some critical challenges remain to be addressed. The first challenge is the single topological representation of drugs. Most deep learning methods—particularly substructure-based methods—represent drugs as molecular graphs with atoms as nodes and bonds as edges. Although this atom-centric topological representation is highly consistent with the real-world molecular structure paradigm, it only defines the adjacency relationships between atoms and neglects a more refined topological description of covalent bonds (e.g., the adjacency relationships between bonds in functional groups), leading to the lack of bond-level semantics in drug representations. It is worth noting that covalent bonds play an extremely important role in the structure-activity relationship of drugs. For example, high-strength covalent bonds enable drugs to resist in vivo degradation mediated by the metabolic enzyme cytochrome P450 (CYP), thus prolonging their half-lives.^19,20^. Therefore, the lack of bond-level semantics would lead to suboptimal representation when predicting DDI. The second challenge that follows is the cognitive bias in model interpretability. Atom-centric methods capture key substructures by visualizing the importance weights of atoms within the drug pair, thus endowing model with interpretability. However, the above process focuses solely on identifying key substructures from the atomic-level semantic perspective and neglects insights of bond-level semantics, which results in model interpretability biased toward atoms. The third challenge is the limited generalization ability. DDI prediction spans a range of tasks, from determining whether an interaction exists between a pair of drugs to quantifying the pharmacokinetic (PK) impact of one drug (the perpetrator) on another drug (the victim). However, current methods narrowly define DDI prediction as a specific task, which separates the relationships between different task types. As a result, the performance advantages of these methods are confined to a single task and cannot generalize to comprehensive DDI prediction.

To address the above challenges, here we propose DualTopoDDI—a dual-topology-enhanced interpretable deep learning model for DDI prediction. By introducing topology inversion as the fundamental technical design, and deriving novel message-passing schemes and other techniques from it, DualTopoDDI captures both atom- and bond-level semantics and delivers interpretable insights from molecular to mechanistic perspectives. As an effort to comprehensively evaluate the risks associated with commonly used medications, we apply DualTopoDDI to predict approximately 9.2 billion DDI entries involving 2,567 approved drugs, the largest in silico evaluations for marketed drugs as far as we are concerned. Among them the high-confidence predictions (confidence score > 0.9) accounts for 97.99%. With regard to interpretation, DualTopoDDI leverages the dual-topology representation so that to recognize the critical atoms and bonds responsible for drug actions. The pinpointed substructures align well with the literature-reported ones. The interpretability of DualTopoDDI also extends to mechanistic level. By training DualTopoDDI to predict particular DDI types between two drugs, we can further decipher mechanisms underlying the complicated adverse effects of multiple drug combinations, such as the controversial cardiotoxicity of COVID-19 medications discussed in this study. Finally, extensive evaluations on 11 benchmark datasets across four types of tasks, also the challenging cold-start scenarios, indicate that DualTopoDDI consistently achieves state-of-the-art (SOTA) or competitive performance against the cutting-edge models, showing excellent generalization ability.

## Results

### Overview of DualTopoDDI

DualTopoDDI converts a drug molecule into a line graph with bonds as nodes and atoms as edges via performing topological inversion on a molecular graph (termed node graph) with atoms as nodes and bonds as edges. In this way, node-line dual-topology drug representations can be constructed to represent single drugs, which replenishes atom- and bond-level semantics. On this basis, we design the Weave Message Passing Neural Network (Weave MPNN), which takes the node-line dual-topology drug representation as input and alternately updates the atoms in the node graph topology and the bonds in the line graph topology, thereby reducing the atom-bond semantic gap caused by topological heterogeneity. Afterwards, we feed the updated node-line dual-topology representations from different Weave MPNN layers into the Cross-topology consensus enhancer. Specifically, different attention coefficients are assigned to different layers of node graph topologies and line graph topologies, thus capturing their respective substructures. In this process, a consistency constraint is imposed between the two topologies to strengthen their substructure consensus, and thus synergistically locate the key substructures or functional groups in combination with atom-bond semantics. Finally, we introduce the Cross-entity interaction module, which integrates the aforementioned substructure knowledge to systematically simulate the atomic- and bond-wise affinities between two drug entities, thereby inferring drug-drug interactions. In this process, atoms or bonds with strong affinities for another drug are identified, endowing the model with dual-topology molecular interpretability. The overall architecture of DualTopoDDI is shown in Figure 1.

**Figure 1.**
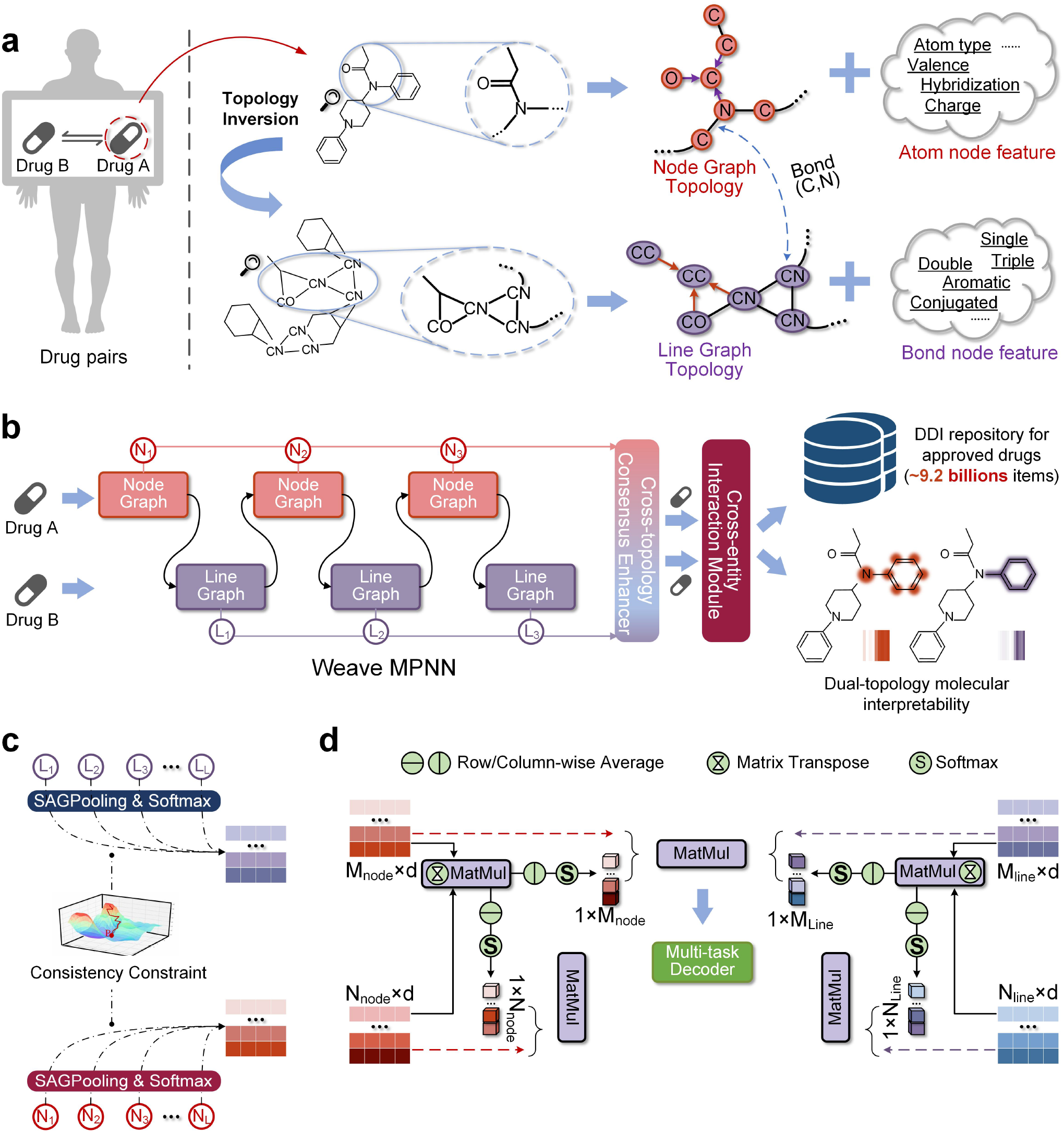
Overview of DualTopoDDI. **a**. The construction of node-line dual-topology drug representation via the operation of topological inversion. **b**. The diagram of DualTopoDDI architecture. **c**. The diagram of cross-topology consensus enhancer. **d**. The architecture of cross-entity interaction module.

### High-throughput prediction of 9.2 billion DDI events for marketed drugs

Polypharmacy is increasingly common in clinical practice, yet systematic experimental characterization of every possible drug pair is neither feasible nor cost-effective. In this section, we first conduct a high-throughput prediction of drug-drug interactions between any two among all marketed drugs to offer a comprehensive toxicity map before they surface in real-world use. In total we offered 9.2 billion DDI prediction results for all possible event types between all 2,567 approved drugs (Supplementary Data 1)^5,6^. Depending on the prediction scenarios, the entries can be classified into three complementary categories.

The first is a standard *binary* view, where a preliminary judgement regarding whether the two drugs have interactions is given. Under this branch, we provide 6.58 million entries with 96.56% of them owing very high confidence (confidence score>0.9). Second, in terms of a *mechanistic* view, the DDI events are classified into 86 predefined mechanistic interaction types, for instances, the increase of anticoagulant activities^21^, the increase of hypocalcemic activities^22^, etc. In this category, we provide 566.47 million interaction results covering all 86 mechanism-driven labels for each drug pair. Despite the challenging nature in recognizing the more detailed interaction types, DualTopoDDI predicts all samples with 98.15% having high confidence. Finally, a more fine-grained interaction type recognition, which contains 8.66 billion *phenotype*-level prediction results, is presented as the third category. It covers myriad real-world phenotypic symptoms like bradycardia and flatulence, containing up to 1,316 predefined phenotype labels. This fine-grained setting challenges the model to capture subtle phenotypic side effect signals. Despite this, DualTopoDDI maintains high confidence for most (97.98%) of its predictions.

To sum up, we employed DualTopoDDI to predict DDIs for 2,567 marketed drugs under various scenarios ranging from basic judgements to very detailed mechanism- and phenotype-level event types. The above three categories together contribute to more than 9.2 billion predictions of DDI events. As far as we have concerned, this is the first and largest prediction effort dedicated to the DDI events of marketed drugs. It is particularly noteworthy that the averaged confidence score of all 9.2 billion predictions is up to 0.99, with high-confidence predictions (>0.9) accounting for 97.99%. As a resource to potentially promote further pharmaceutical innovations, all these predictions are publicly available via the online data repository.

### Dual-topology enables molecular-level interpretation of DDI mechanisms

A key strength of DualTopoDDI is its ability to leverage dual-topology representations to deliver molecular-level interpretability of DDI events. By modeling both atom-centric and bond-centric views, DualTopoDDI pinpoints the specific substructures responsible for drug actions. In this section, we demonstrate the interpretability of DualTopoDDI using an enzyme inhibition dataset (Supplementary Data 2). Specifically, the enzyme inhibition depicts a particular DDI scenario where a drug (termed perpetrator) affects the pharmacokinetic of another drug (termed victim) by inhibiting the relevant metabolic enzymes^23^. In this context, rapid identification the key substructures in the perpetrator drug is vital for explaining the DDI mechanism^24^. Accordingly, we collected 58 available perpetrator drugs (Supplementary Data 3) with well-characterized key substructures that literature reports as responsible for metabolic enzyme inhibition. These molecules and in particular their substructures act as ground truth to validate the molecular-level interpretability of models. Next, we retrieved a total of 16,049 positive DDI events (Supplementary Data 2) involving the 58 drug molecules as perpetrators from existing datasets or via the mediated enzyme. For all these interactions, DualTopoDDI was able to identify 10,587 entries that cover 57 drug molecules (Supplementary Data 4). Leveraging the dual-topology representations in DualTopoDDI, we projected the atom- and bond-wise attention scores onto the molecular structure of each perpetrator to highlight the model-identified key atoms and covalent bonds. The attention scores were achieved by summing the atom- and bond-wise attention scores across all DDIs related to a perpetrator and then scaling them to [0, 1] across atoms or bonds. The visualization of key atoms and key bonds are shown in Figure 2a and b, respectively.

**Figure 2.**
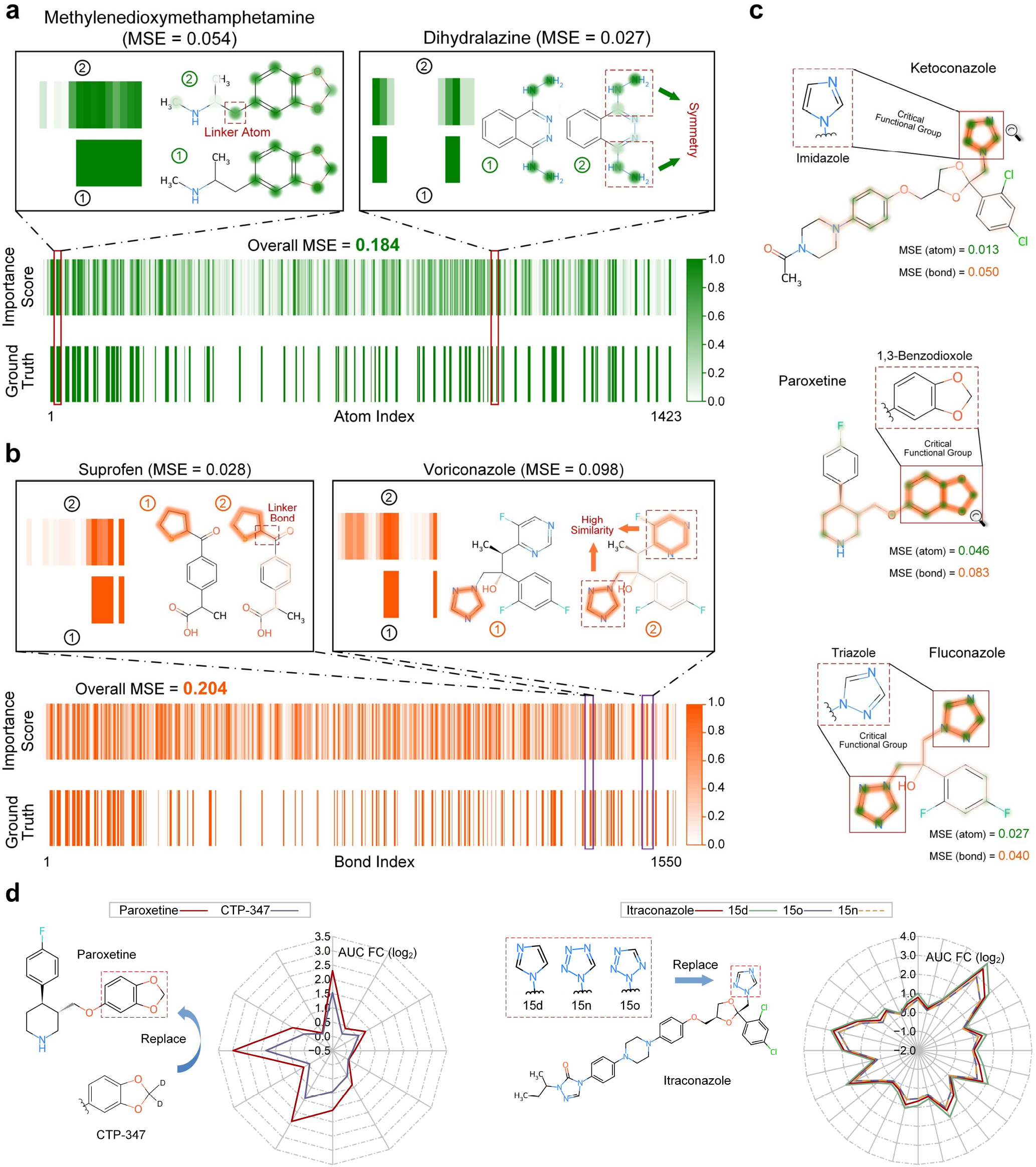
Comprehensive case studies of substructural insight with DualTopoDDI. **a**. The atom-wise visualization of key substructures via dual-topology molecular interpretability. **b**. The bond-wise visualization of key substructures via dual-topology molecular interpretability. **c**. Visualization case analyses of Ketoconazole, Paroxetine, and Fluconazole using combined atom- and bond-wise importance scores. **d**. Case analyses of paroxetine and Itraconazole for structural modifications.

According to Figure 2, the atom-wise and bond-wise importance scores assigned by DualTopoDDI exhibit a high degree of overlap with the literature-reported key substructures, with overall MSE values of 0.184 and 0.204, respectively (Supplementary Data 5 and 6). In particular, for the atom-centric topology, we highlight two exemplar drugs with small MSEs, namely methylenedioxymethamphetamine (MSE=0.054) and dihydralazine (MSE=0.027). As shown in the enlarged panel in Figure 2a, the model-inferred critical atoms align well with the ground-truth key substructures known to affect metabolic enzymes. Additionally, the model also allocated a high attention score to the linker atoms connecting these substructures. This observation further demonstrates DualTopoDDI’s insight into substructures or functional groups, and suggests that linker atoms may also play an indispensable role in the induction or inhibition of metabolic enzymes. Another interesting observation is that, for the symmetric key substructure in dihydralazine, DualTopoDDI assigned consistent importance scores, which underscores DualTopoDDIs ability in understanding and grasping the underlying structural motifs in molecules.

Similarly, in terms of the bond-centric topology, suprofen and voriconazole are illustrated, which yielded MSEs of 0.028 and 0.098, respectively (Supplementary Data 6). As shown in the enlarged view in Figure 2b, in addition to the precise capture of the bonds in the key substructures, the model likewise pinpointed the linker bond, mirroring its identification of the linker atoms. Another finding from voriconazole is that, DualTopoDDI accurately captured the known key substructure and, interestingly, also identified 5-fluoropyrimidine—a highly similar functional group to the known one. This underscores DualTopoDDI’s potential to uncover novel DDI mechanisms.

Finally, the two topologies in DualTopoDDI may also reveal critical substructures synergistically. In this aspect, three drugs namely ketoconazole, paroxetine, and fluconazole are visualized, as shown in Figure 2c. It can be observed that the model-identified key atoms and covalent bonds have markedly higher importance scores compared to other atoms and chemical bonds. These key elements together form the complete substructures or functional groups reported in the literature, further validating DualTopoDDI’s good interpretability of DDI mechanisms at the molecular level.

### Computation-driven structural modification to attenuate metabolic DDIs

In the de novo drug design, structural modifications of candidate molecules are frequently implemented to achieve desired physicochemical properties^25,26^. Identically, researchers attempt to optimize key substructures of perpetrator drugs to retain therapeutic efficacy while alleviating their inhibitory effects on metabolic enzymes, aiming to mitigate adverse reactions caused by the exposure of victim drugs. Given the immense chemical space, it is infeasible to systematically evaluate all possible structural modifications through *in vivo* or *in vitro* experimentations. In this regard, DualTopoDDI offers a computation-driven alternative for rapidly screening effective structural modifications. Using paroxetine as a case study—a potent, selective serotonin reuptake inhibitor (SSRI) for the treatment of depression, we collected 12 paroxetine-perpetrated DDIs (Supplementary Data 7). In particular, the 1,3-benzodioxole group in paroxetine leads to quasi-irreversible inactivation of the metabolic enzyme CYP2D6, thereby elevating plasma concentrations of co-administrated victim drugs^27^. By contrast, deuterated paroxetine (CTP-347), which replaces protium with deuterium at the methylenedioxy carbon of the 1,3-benzodioxole structure, demonstrates attenuated CYP2D6 inhibition^28,29^. Accordingly, the plasma concentrations of victim drugs decrease. To validate whether DualTopoDDI can recapitulate this mechanism, we employed DualTopoDDI to calculate concentration changes of 12 victim drugs when paired with paroxetine and the deuterated analogue CTP-347, respectively. The concentration change is reported as AUC FC (i.e., area under the plasma time-concentration curve fold change, see Methods for more details), and a higher AUC FC value indicates a more severe metabolic DDI. As illustrated in Figure 2d, CTP-347 universally reduced the AUC FC across all victim drugs. Particularly for eliglustat (DB09039)—a therapeutic agent for Gaucher disease type 1^30^—the log2(AUC FC) decreased from 2.98 to 1.83.

Similarly, itraconazole—a triazole broad-spectrum antifungal agent—exerts its antimicrobial effects by inhibiting the biosynthesis of ergosterol in fungal cell membranes^31^. Recent studies have revealed its significant antitumor potential, particularly through antiangiogenic activity and modulation of oncogenic signaling pathways. However, its inhibition of CYP3A4 increases the plasma concentration of victim drugs, thereby constraining its clinical application. In this regard, structural modification of the 1,2,4-triazole ring group in itraconazole offers a promising solution^31^. Similar to the case study of paroxetine, we collected 28 drug pairs where itraconazole acts as the perpetrator. We first calculated AUC FC values of all victim drugs when accompanied with the original itraconazole molecule as a control group. Next, we obtained three itraconazole analogues (i.e., 15d, 15n, and 15o) by replacing the 1,2,4-triazole with imidazole, 1-tetrazole-yl, and 2-tetrazole-yl groups, respectively. Afterwards, we re-calculated the AUC FC values of the 28 victim drugs by substituting itraconazole with these analogs (Supplementary Data 7). As shown in Figure 2d, analogs 15n and 15o display an overall downward trend in the AUC FC of victim drugs, indicating attenuated CYP3A4 inhibition compared to the control group. In contrast, 15d exhibits an increased AUC FC. These observations align well with experimental evidence^31^.

### DualTopoDDI uncovers cardiac risks of controversial COVID-19 drug combinations

The rapid repurposing of COVID-19 therapies often outpaced rigorous safety evaluation, leading to lingering debate over the arrhythmogenic potential of certain drug combinations. At the time, traditional invitro and clinical investigations were slow and frequently produced conflicting evidence on their cardiac risks. To address this retrospective gap, we leverage DualTopoDDI to predict and mechanistically interpret the cardiotoxicity of two high-profile COVID-19 regimens.

We first investigated a controversial combination, i.e., hydroxychloroquine and azithromycin. Hydroxychloroquine was rapidly repurposed early in the COVID-19 pandemic and frequently co-administered with the macrolide antibiotic azithromycin to augment antiviral activity. Both hydroxychloroquine and azithromycin block the hERG (KCNH2) potassium channel, prolonging the QT interval and raising the likelihood of Torsade de Pointes arrhythmias in combination^32,33^. However, subsequent clinical studies have reported conflicting evidence on whether the combination of hydroxychloroquine and azithromycin actually induces malignant ventricular arrhythmias^34,35^. Here, we apply DualTopoDDI to this controversial regimen—predicting its cardiac risks and pinpointing the molecular features driving potential pro arrhythmic interactions. According to the predictions, DualTopoDDI ruled out with very high confidence (>0.99) that the two drugs increase the cardiotoxic activities of each other. Using more fine-grained interaction type labels, DualTopoDDI further denied that the two drugs synergistically result in arrhythmogenic activities or mild rhythm changes like bradycardia and tachycardia. All these predictions have very high confidence scores. These observations align well with compelling real-world clinical evidence^34-36^, which are considered golden standards. Interestingly, the model suggested that the risk of QT-interval prolongation can be increased upon combination (confidence score=0.958), which has been reported in animal experiments and clinical observations^33^. QT interval prolongation is generally viewed as a warning sign, since it can precipitate more serious arrhythmias such as Torsade de Pointes. However, clinical observations have shown that the QT prolongation caused by the hydroxychloroquine-azithromycin combination is mild and has not led to severe, life-threatening arrhythmic events^36,37^. DualTopoDDI partially explains the controversy surrounding its cardiac safety by mechanistically predicting QT-interval prolongation while ruling out malignant arrhythmic activities and other cardiotoxic events.

The next high-profile drug combination investigated here is the Paxlovid and quinidine. Paxlovid is a two-component antiviral therapy combining nirmatrelvir (a SARS CoV 23CL protease inhibitor) with ritonavir, a pharmacokinetic booster whose primary role is potent CYP3A4 inhibition. While nirmatrelvir targets viral replication, ritonavirs suppression of CYP3A4 can dramatically increase plasma levels of co-administered drugs thus leading to well-documented cardiotoxicity^38^. Quinidine, a common antiarrhythmic agent, is among these cardiotoxic drugs upon co-administration. Here we show that DualTopoDDI is capable of mechanistically interpreting the high cardiac risk of the Paxlovid-quinidine combination. We first employed DualTopoDDI to predict the cardiotoxicity of nirmatrelvir and quinidine. In this case, DualTopoDDI showed that nirmatrelvir neither increases cardiotoxic activities nor arrhythmogenic activities—all with very high confidence (>0.99)—suggesting that nirmatrelvir, as the antiviral component in Paxlovid, is not associated with the adverse cardiac events. We then tested the combination of ritonavir and quinidine. DualTopoDDI confidently predicted that ritonavir increases the arrhythmogenic activities of quinidine, which aligns well with existing evidence. More importantly, we found that DualTopoDDI is capable of capturing the most critical mechanism underlying the increased risks of arrhythmias. Specifically, DualTopoDDI predicted that the serum concentration of quinidine can be increased when combined with ritonavir (confidence score>0.99), but not vice versa (confidence score>0.99). This suggests that the model is able to differentiate the causal relationship, recognizing that ritonavir can increase quinidine concentration, while quinidine does not affect ritonavir. Another similar result also proves this, where DualTopoDDI predicted that the metabolism of quinidine can be decreased when combined with ritonavir but not the other way around.

To further demonstrate the arrhythmogenic mechanisms of this combination, we built an anatomically detailed virtual heart model (Figure 3) and incorporated the drug-induced modulation of cardiac ion channels into the virtual heart. We then conducted in-silico experiments to demonstrate how the drug combination can induce malignant arrhythmias. The simulation results are shown in Figure 3. It can be observed that the heart with quinidine alone generated a normal and regular morphology on pseudo-ECG, reflecting a sinus rhythm without any arrhythmogenic activities. Further visualization of electrophysiological activities in the virtual heart demonstrated that the excitation waves propagated regularly (Figure 3a), suggesting that quinidine alone did not induce apparent pathological changes. This observation is attributed to the fact that a low-dose of quinidine does not lead to arrhythmias. However, the situation was changed when quinidine was administrated in combination with ritonavir. As shown in Figure 3b, due to the inhibition of quinidine metabolism by ritonavir, the serum concentration of quinidine was kept at a high level, and the regular Q-T pattern swiftly evolved into ventricular arrhythmias, manifesting as chaotic waves on the pseudo-ECG. According to a further visualization, such event initiated with a sudden ectopic activity oriented from the bottom of the ventricle, which then evolved into fibrillation-like excitation waves across the whole ventricle. The simulation results proved the proarrhythmic effects of the combination of quinidine and ritonavir. These results together validated the powerful ability of DualTopoDDI not only in recognizing general DDI types such as cardiotoxicity, but also in deciphering the complicated underlying mechanisms like the inhibited metabolism and the increased serum concentration, particularly in scenarios of multiple drug interactions.

**Figure 3.**
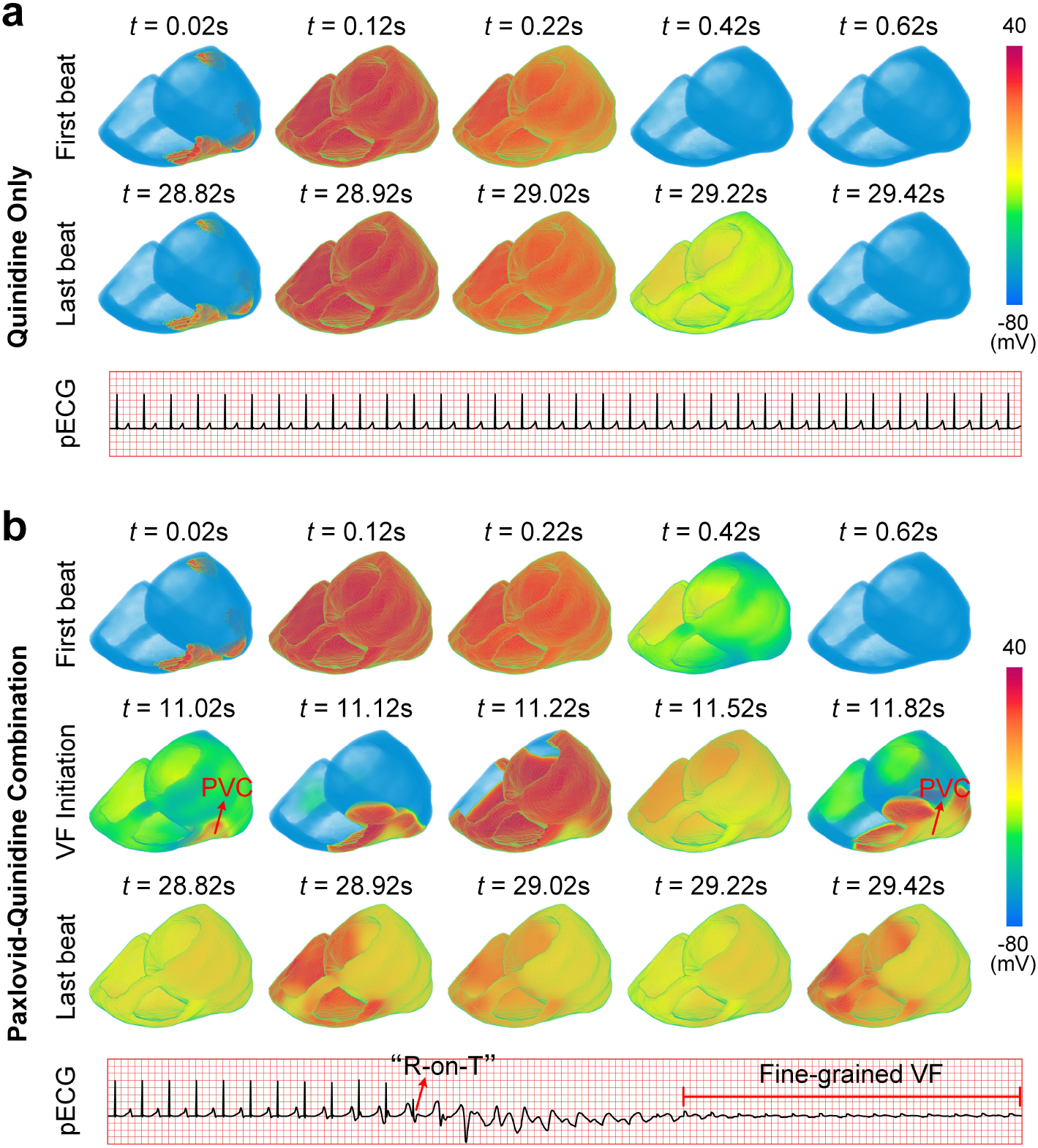
Simulations of the effects of COVID-19 drug combinations using a virtual heart. **a**. Applying quinidine only as a control group, where no obvious arrhythmias were observed. This observation show that quinidine alone does not induce cardiotoxicity. **b**. Applying quinidine together with Paxlovid. It can be observed that ventricular arrhythmias initiated with a premature ventricular contract (PVC) at around 11.02 s, which later evolved into fine-grained ventricular fibrillations, suggesting a potential cardiotoxicity of Paxlovid-Quinidine combinations.

### DualTopoDDI delivers consistent performance across various DDI scenarios

We benchmarked the performance of DualTopoDDI against existing methods on 11 datasets that cover various DDI scenarios. From a computational perspective, the 11 datasets used in this work can be categorized into four types DDI prediction tasks (see Methods), including determining the presence of interactions between drug pairs (standard DDI task), identifying specific interaction types (DDI type prediction task), predicting drug combination effects on PK (metabolic DDI classification task), and quantifying the magnitude of PK alterations (PK fold change prediction task). In this section, we compared the performance of DualTopoDDI with multiple baseline methods on the above four types of DDI benchmark tasks to evaluate the generalization ability of the proposed model.

For the standard DDI task, it can be seen from Figure 4 that DualTopoDDI achieved optimal performance across all metrics on the DeepDDI and ZhangDDI datasets, except for Rec (suboptimal). Notably, on the DeepDDI dataset, DualTopoDDI attained near-perfect scores (>99) in nearly all metrics. On the other hand, while DualTopoDDI exhibited slightly lower performance on the ChChMiner dataset, it still maintained competitive results compared to the top-performing method MeTDDI, i.e., 99.22 vs. 99.79 (AUC), 99.66 vs. 99.97 (Prec), 99.89 vs. 99.97 (AP).

**Figure 4.**
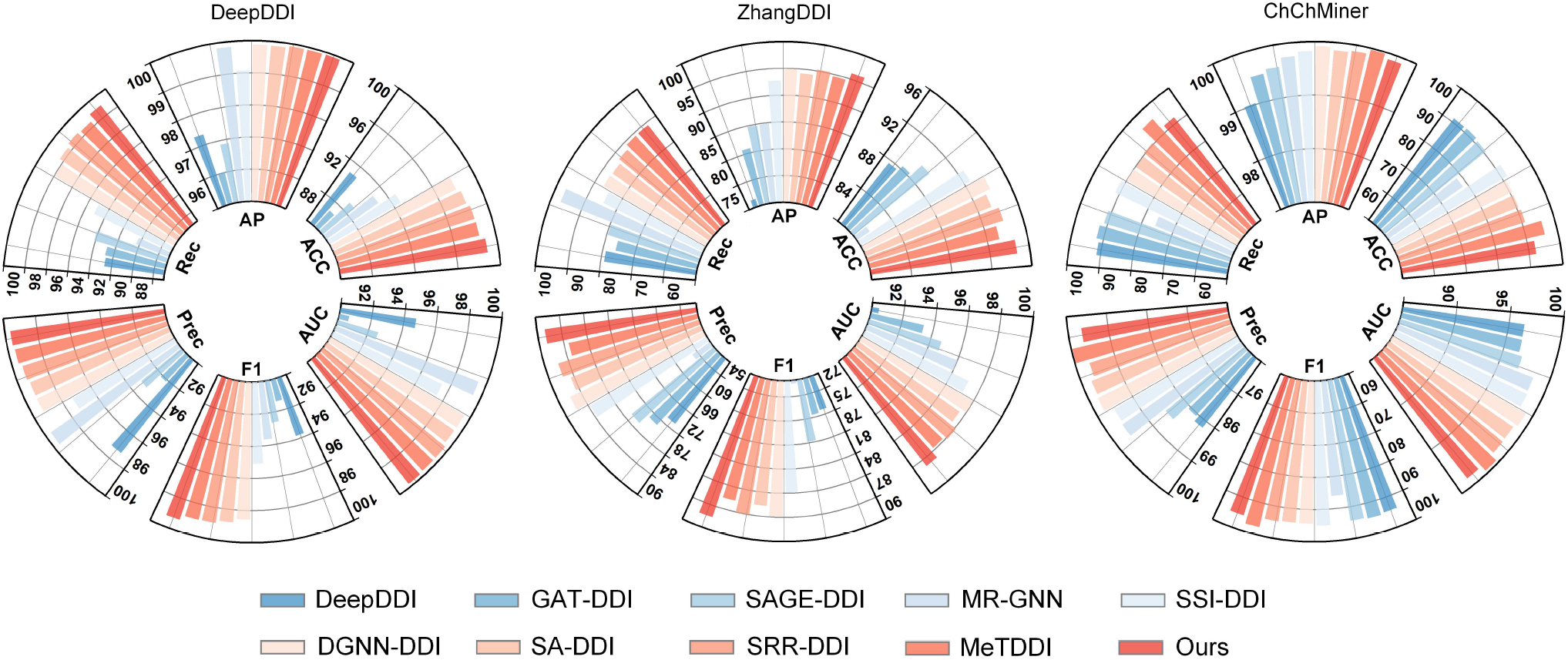
Performance comparison of DualTopoDDI with baseline methods on three datasets of standard DDI task, namely DeepDDI, ZhangDDI, and ChChMiner.

For the DDI type prediction task, DualTopoDDI also demonstrates its superiority, as shown in Figure 5. Specifically, on the DrugBank dataset, DualTopoDDI immensely outperforms the suboptimal method SRR-DDI, achieving optimal performance across metrics. Especially for ACC and F1 metrics, DualTopoDDI is the only method to surpass the 97, i.e., ACC = 97.51 and F1 = 97.54. Similarly, DualTopoDDI attains optimal results in all metrics except the suboptimal performance on the Prec metric of the TWOSIDES dataset (i.e., 82.74 vs. 83.06). On this basis, DualTopoDDI was further evaluated on the cold-start datasets DrugBank-S1 and DrugBank-S2, where the model achieved optimal performance in terms of ACC, AUC, Prec, and MSE performance of DualTopoDDI and other methods on AUC_FC and AUC_FC_External datasets.

**Figure 5.**
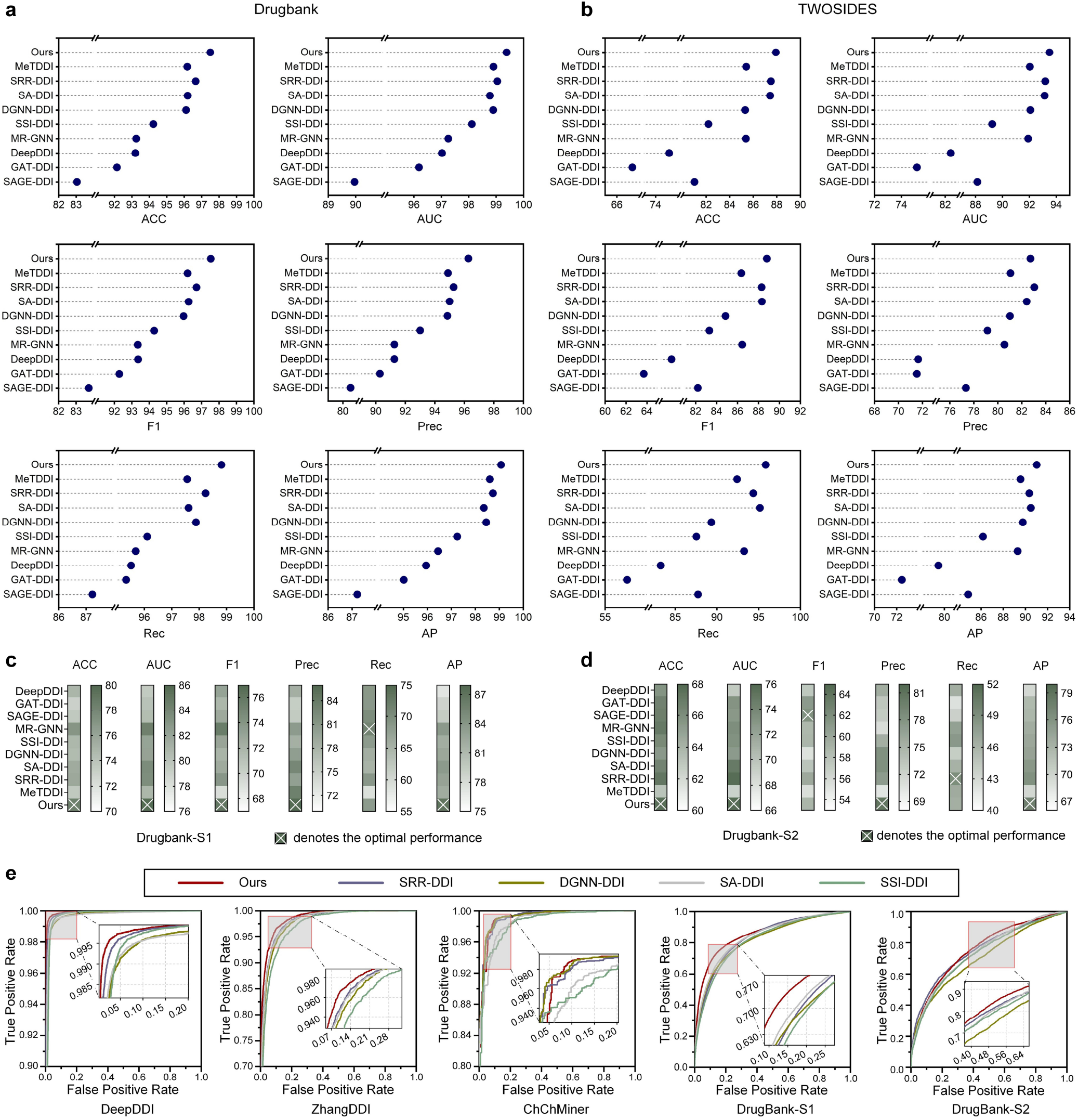
Performance comparison of DualTopoDDI with baseline methods on the DDI type prediction task. **a**. Performance of DualTopoDDI and baseline methods on the DrugBank dataset. **b**. Performance of DualTopoDDI and baseline methods on the TWOSIDES dataset. **c**. Performance of DualTopoDDI and baseline methods on the DrugBank-S1 dataset. **d**. Performance of DualTopoDDI and baseline methods on the DrugBank-S2 dataset. **e**. ROC curves of DualTopoDDI and representative cutting-edge methods.

AP metrics. These results demonstrate its robustness in predicting DDIs for novel drugs. Furthermore, the ROC curves of DualTopoDDI on the above datasets are shown in Figure 5e, which further demonstrates the superiority of DualTopoDDI.

For both the metabolic DDI classification task and PK fold change prediction task, DualTopoDDI identically achieved impressive performance across all metrics on each dataset (Figure 6), including external test sets, as anticipated. Particularly for the MMDDI dataset, DualTopoDDI outperformed the suboptimal method DGNN-DDI by 2.63% on ACC and 2.99% on F1, i.e., 97.90 vs. 95.27 (ACC) and 97.86 vs. 94.87 (F1). Additionally, on the AUC_FC_External, DualTopoDDI demonstrated overwhelming advantages compared to the suboptimal method MeTDDI, with a substantial margin of 0.15 in term of the MSE metric. Details of performance comparisons can be found in Supplementary Tables 1-4. To visually demonstrate DualTopoDDI’s superiority in the PK fold change prediction task, we collected ground-truth labels and corresponding DualTopoDDIs prediction from the AUC_FC and AUC_FC_External datasets, so that scatter plots can be generated, as shown in Figure 6e. The fitted function derived from the scatter distribution closely aligns with the diagonal line, further validating DualTopoDDIs robustness to quantify PK alterations. On this basis, we statistically visualized the MSE values for each test sample across both datasets using different methods (Figure 6f). While MSE values from all methods are concentrated near zero, DualTopoDDI obviously exhibits a shorter-tailed distribution.

**Figure 6.**
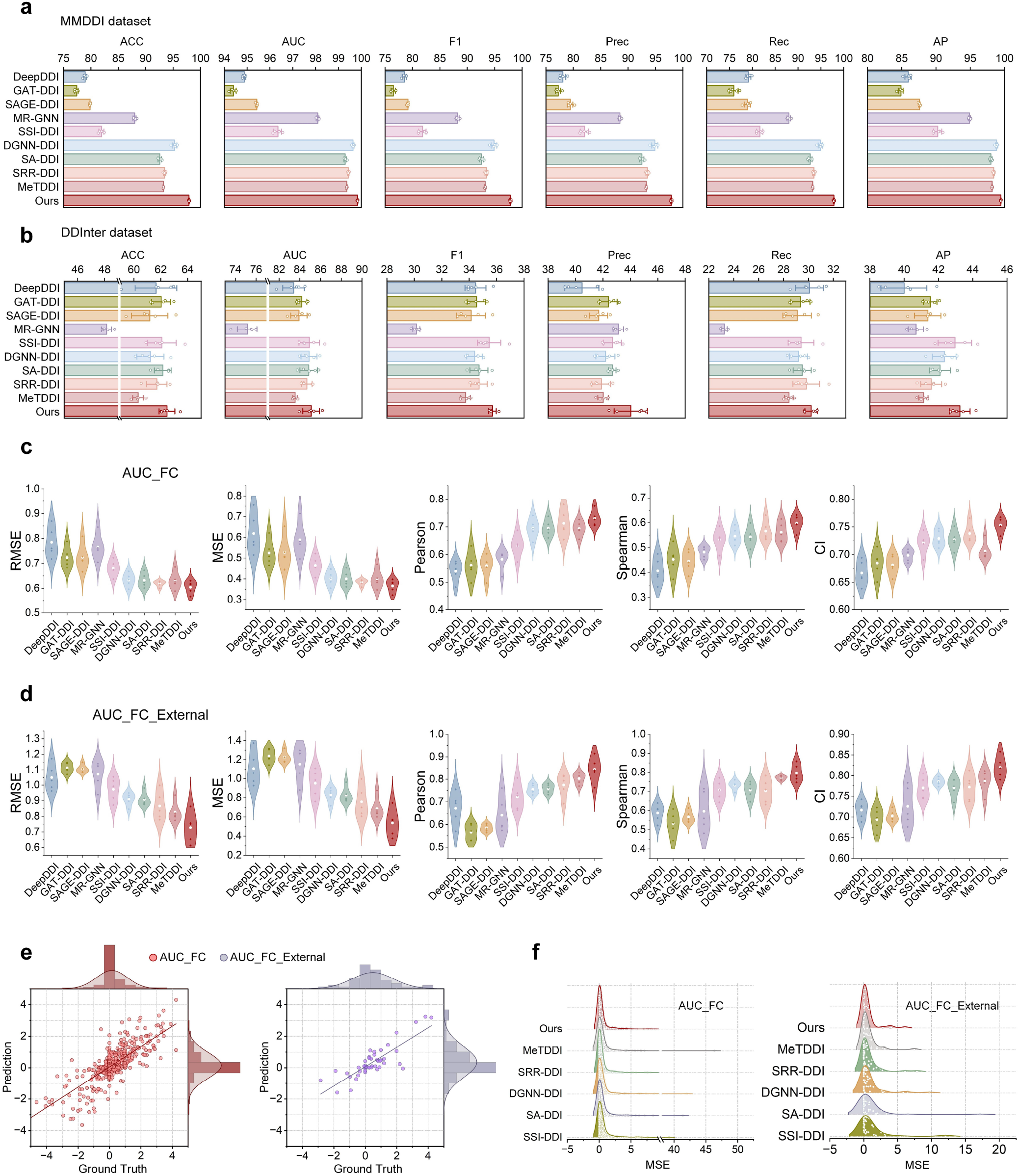
Performance comparison of DualTopoDDI with baseline methods on the metabolic DDI classification task and PK fold change prediction task. **a**. Performance of DualTopoDDI and baseline methods on the MMDDI dataset. **b**. Performance of DualTopoDDI and baseline methods on the DDInter dataset. **c**. Performance of DualTopoDDI and baseline methods on the AUC_FC dataset. **d**. Performance of DualTopoDDI and baseline methods on the AUC_FC_External dataset. **e**. The predicted values and corresponding ground truth of DualTopoDDI on AUC_FC and AUC_FC_External datasets. **f**. MSE performance of DualTopoDDI and other methods on AUC_FC and AUC_FC_External datasets.

### In-depth model architecture analysis from a computational perspective

We first performed dimensionality reduction via the t-SNE algorithm^39^ on drug pair representations from the hidden layer of DualTopoDDI across diverse datasets, visualizing them in a two-dimensional feature space (Figure 7a). The distinct clustering patterns observed demonstrate DualTopoDDIs ability to learn discriminative drug pair embeddings, thus reflecting the superiority of DualTopoDDI on DDI task. Furthermore, to validate the contributions of individual modules in DualTopoDDI to DDI prediction performance, we also conducted systematic ablation experiments in this section. Specifically, we sequentially removed each module to generate different model variants, which were subsequently retrained and evaluated on the DrugBank dataset. The resulting model variants were defined as follows:

**Figure 7.**
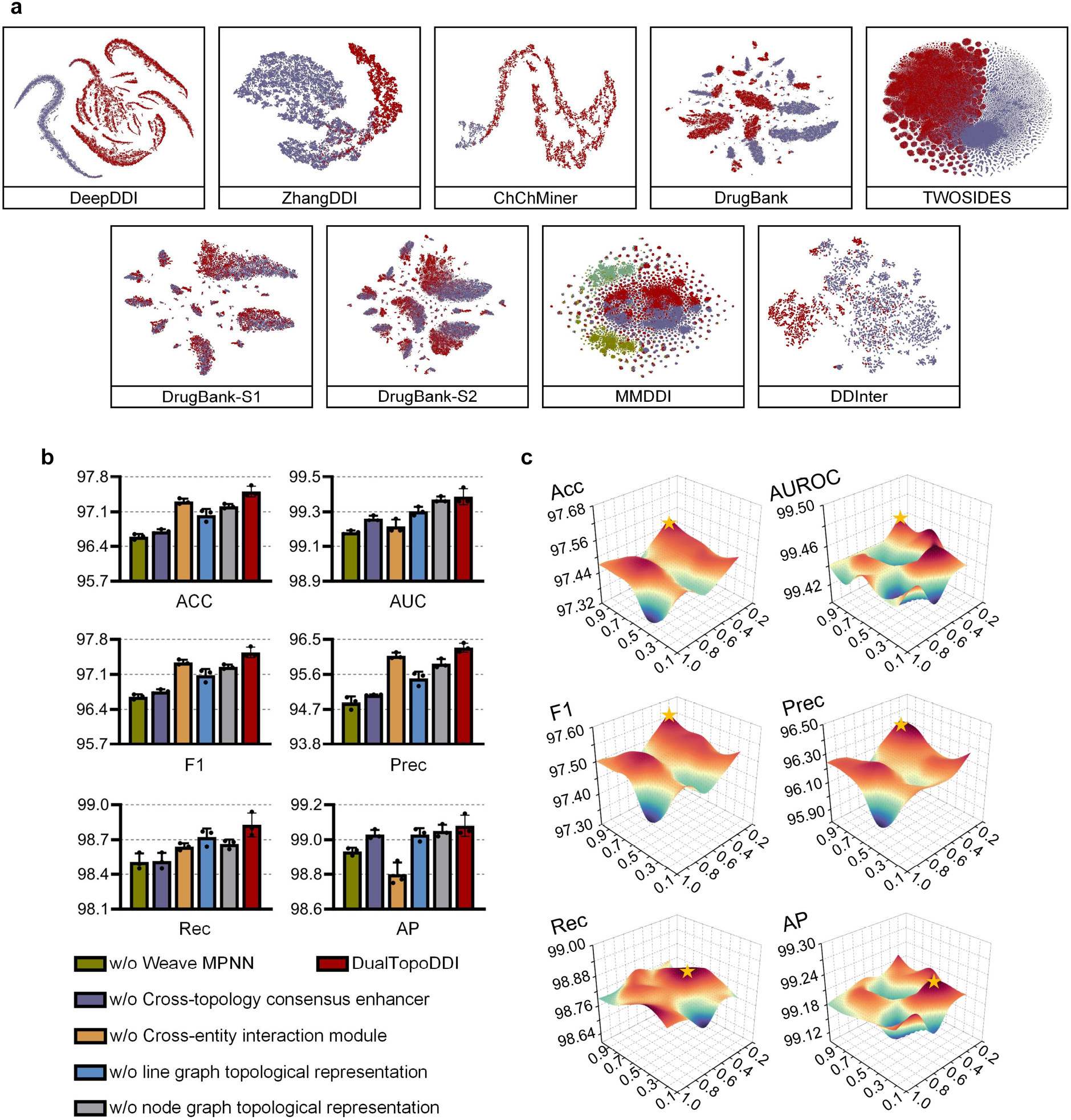
The analysis results of DualTopoDDI architecture. **a**. Distribution of sample points (drug pairs) on different datasets under two-dimensional feature space. **b**. Ablation results of DualTopoDDI on the DrugBank dataset. **c**. Hyperparameter optimization of λ_1_ and λ_2_ in DualTopoDDI.

1. w/o Weave MPNN: In accordance with conventional operation, each Weave MPNN layer was replaced with two independent graph convolutional layers dedicated to feature extraction from node graph and line graph representations of drug molecules, respectively.
2. w/o cross-topology consensus enhancer: The output of Weave MPNN was directly fed into the cross-entity interaction module and the cross-topology consensus enhancer is no longer considered.
3. w/o cross-entity interaction module: Similarly, a pair of drug representations was directly input to the MLP decoder without going through the cross-entity interaction module for subsequent DDI prediction.
4. w/o node-line dual-topology drug representation: Only the node graph topological representations of drugs were retained, which entails that line graph topological representations—and the underlying bond-level semantic—were excluded from consideration.

Figure 7b presents the detailed ablation results, which reveals that removing any module from the intact DualTopoDDI led to performance degradation, thereby substantiating the indispensability of each module. Among them, DualTopoDDI without Weave MPNN (w/o Weave MPNN) obtained the worst performance overall, particularly evident in the ACC and Prec metrics, i.e., ACC=96.59 and Prec=94.87. In contrast, the performance degradation observed in the w/o dual-topology drug representation variant is notably less severe than that of w/o Weave MPNN. This suggests that while the latter incorporated bond-level semantics, its inability to resolve atom-bond semantic gap arising from topological heterogeneity led to conflicts between bond- and atom-level semantics, resulting in further performance degradation. Afterwards, we optimized some critical hyperparameters in DualTopoDDI, including coefficients λ_1_ and λ_2_ of different loss terms, Weave MPNN layers, the number of heads in cross-topology consensus enhancer, and different calculation operations of atom/bond-wise affinity between drugs. As shown in Figure 7c, DualTopoDDI attains relatively superior performance on metrics excluding Rec and AP when λ_1_ approaches 1 and λ_2_ approaches 0.4. Regarding Rec and AP metrics, setting λ_2_ to 0.4 remains the optimal choice. Similarly, it can be seen from Figure 8 that as the number of Weave MPNN layers increases, DualTopoDDIs overall performance exhibits an upward trend (excluding the Rec and AP metrics). Notably, when the number of Weave MPNN layers is set to 2, the model performance degrades drastically. On the other hand, according to DualTopoDDIs performance, we ultimately set the number of attention heads in cross-topology consensus enhancer to 2 and adopted transpose multiplication to compute atom/bond-wise affinity between drugs.

**Figure 8.**
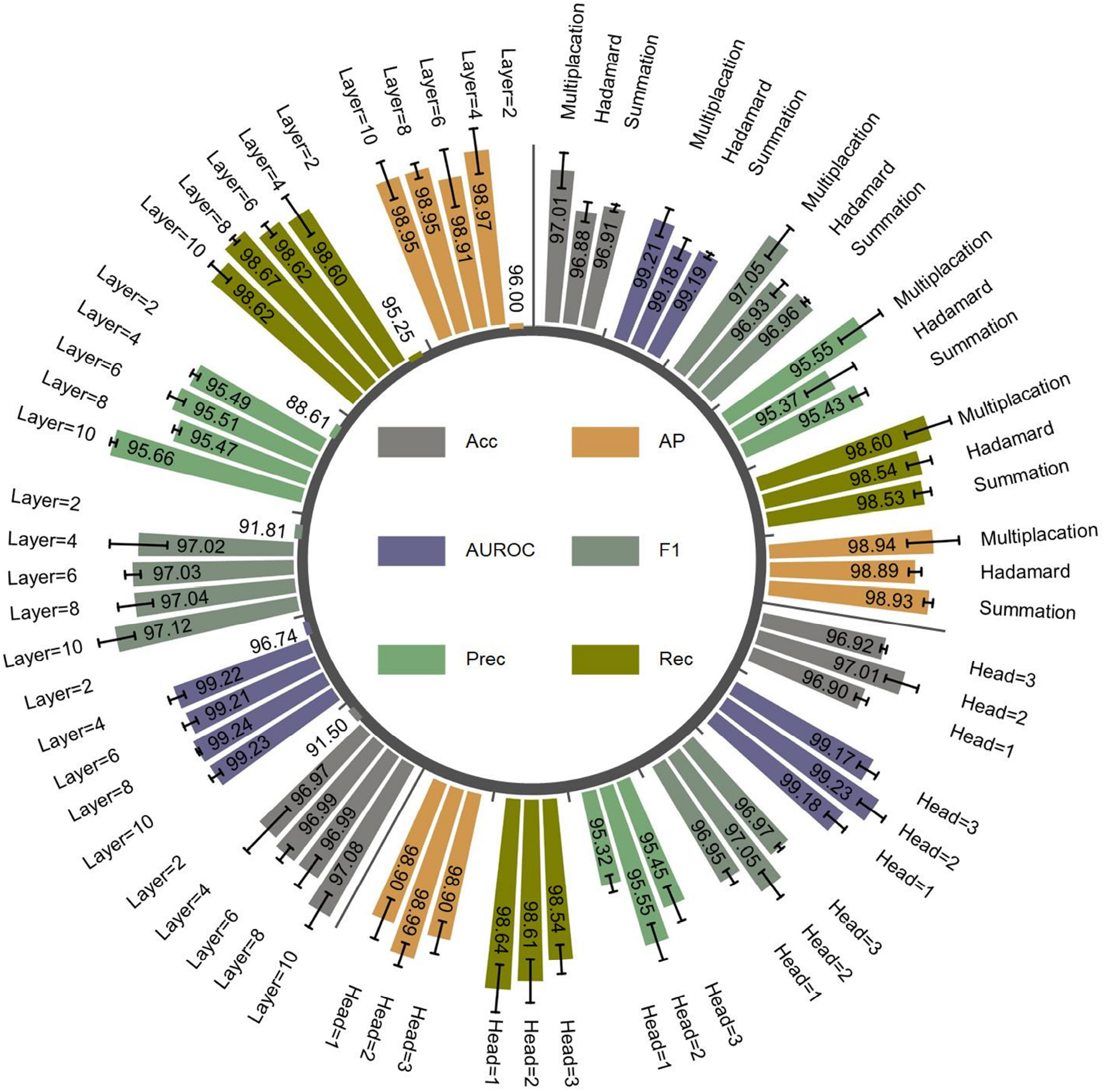
Optimization results of other hyperparameters in DualTopoDDI, including Weave MPNN layers, the number of heads in cross-topology consensus enhancer, and different calculation operations of atom/bond-wise affinity between drugs.

## Discussion

Predicting drug interactions is critical for accelerating pharmacological research and ensuring patient safety. However, existing DDI prediction methods face three critical challenges: (1) single topological representation of drugs, (2) cognitive biases in model interpretability, and (3) limited DDI generalization capability. In this work, we propose DualTopoDDI, an innovative model that captures atom-level and bond-level semantics by constructing node-line dual topological representations for drug molecules. This design not only enhances the models predictive performance on DDI tasks but also provides interpretable insights from the microscopic molecular level (e.g., key substructures driving interactions) to the macroscopic mechanistic level (e.g., controversial cardiotoxicity of COVID-19 medications). Furthermore, DualTopoDDIs performance on 11 benchmark datasets demonstrates its excellent generalization capability across different DDI prediction tasks. On this basis, we also predicted approximately 9.2 billion DDI entries involving 2,567 approved drugs, with high-confidence predictions accounting for 97.99%, thus providing critical decision-making support information for enhancing clinical medication safety and mitigating potential drug-related risks.

On the other hand, as mentioned above, a core strength of DualTopoDDI lies in its interpretability from the microscopic molecular level to the macroscopic mechanistic level. At the microscopic level, DualTopoDDI is capable of identifying crucial atoms and bonds from the drug’s node graph and line graph topologies, respectively, and thus accurately capturing the key substructures driving DDIs (e.g., 1,3-benzodioxole in paroxetine and the triazole ring in fluconazole). The attention scores also demonstrate strong alignment with the established evidence reported in the literature. This microscopic interpretability represents an obvious advancement compared to most DDI methods that focus solely on binary prediction (presence/absence of DDI), as making predictions only about whether DDIs are present, without elucidating why they occurs, severely limits their utility in drug development. Taking paroxetine as an example, DualTopoDDI precisely identified the key substructure within paroxetine responsible for inhibiting CYP2D6, namely the 1,3-benzodioxole (Figure 2c). In this way, the adverse DDI caused by paroxetine can be mitigated by modifying 1,3-benzodioxole while preserving the therapeutic effect. It is worth noting that DualTopoDDI, with microscopic molecular-level interpretability, remains capable of computationally guiding the modifications. Specifically, DualTopoDDI enables rapid prediction of the impact on DDI severity (i.e., AUC FC values of victim drugs) resulting from modifications to key substructures, both before and after the modification. The value of this capability is further exemplified in the paroxetine case study: DualTopoDDI computationally correctly reproduced the phenomenon where the modified paroxetine (CTP-347) exhibited reduced CYP2D6 inhibition, thereby attenuating the adverse DDI. Similarly, for the modification of itraconazole, DualTopoDDI yielded results consistent with clinical trial data (Figure 2d), demonstrating its ability to select effective structural modifications and highlighting its distinctive role in intelligent drug design.

In addition to the interpretability at molecular level, DualTopoDDI also demonstrated its interpretability at the macroscopic mechanism level, as evidenced here through its analysis of cardiotoxicity in COVID-19 drug combinations. Specifically, for hydroxychloroquine and azithromycin, DualTopoDDI confidently refuted any synergistic effect between the drugs regarding pro-arrhythmic activity. Conversely, it accurately identified their propensity to prolong the QT interval. This prediction aligns closely with clinical observations showing that this combination induces only mild QT prolongation and rarely progresses to life-threatening arrhythmias. Similarly, for the Paxlovid and quinidine, DualTopoDDI elucidates the pharmacological mechanism underlying their combination-induced cardiotoxicity from a computational perspective. Specifically, DualTopoDDI not only identified that ritonavir—an antiviral component of Paxlovid—elevated quinidine plasma concentrations by inhibiting CYP3A4, thereby triggering cardiotoxicity, but also established the unidirectional nature of this metabolic inhibition. This mechanistic insight was dynamically validated through a virtual heart model: elevated quinidine concentrations initiated a pathological cascade progressing from localized ectopic activity to fulminant ventricular fibrillation. Critically, DualTopoDDI concurrently excluded any cardiotoxic contribution from nirmatrelvir (Paxlovids another antiviral component), highlighting its precision in causal attribution. This macroscopic mechanistic elucidation, from molecular metabolic disruption (CYP inhibition) to cellular electrophysiological disturbances (ion channel imbalance), and finally to organ-scale arrhythmia, transcends the limitations of traditional DDI prediction models.

## Methods

### Benchmark datasets

In this work, 11 widely used benchmark datasets were utilized to evaluate the performance of our model on 4 types of DDI tasks. For the standard DDI task of determining whether interactions exist, it can be formulated as 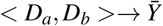. Among them, *D*_*a*_, *D*_*b*_ denote a pair of drugs and 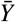 is the prediction score ranging from 0 to 1. In this regard, the ZhangDDI^40^, ChChMiner^41^, and DeepDDI^42^ datasets were introduced. After removing the anomalous data caused by irregular SMILES strings, the three datasets contained 544, 997, and 1704 drugs, respectively, with 45,720, 21,486, and 191,870 positive DDI samples, while negative samples were randomly sampled from non-existent DDIs. Then, the datasets were split into the training set and the test set according to the ratio of 4:1. Additionally, 25% of the training data was randomly selected to form the validation set.

Similarly, the DrugBank^5^ and TWOSIDES^43^ datasets were employed to evaluate the performance of model on the DDI type prediction task, which can be defined as 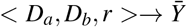. Among them, *r* indicates a specific DDI type. On this basis, DrugBank dataset contained 1,706 drugs, involving 191,808 DDIs and 86 interaction types, each of which described metabolic impacts (e.g., the increase of anticoagulant activities) induced by drug combinations. The other comprised 645 drugs, involving 4,576,287 DDIs and 1,316 interaction types, where each type indicated potential phenotypic symptoms (e.g., bradycardia, flatulence) arising from drug combinations. Negative samples for both datasets were generated following the strategy proposed by Wang et al^44^. Training and test sets were randomly divided at a 4:1 ratio. To further assess model capability in predicting interactions between novel drugs, we utilized two cold-start datasets derived from DrugBank, i.e., DrugBank-S1 and DrugBank-S2^5^. Specifically, 20% of drugs in the DrugBank dataset were regarded as novel drugs while the rest served as known drugs. On this basis, both DrugBank-S1 and DrugBank-S2 share identical training sets containing only known drugs. As for the test set, each drug pair in Drugbank-S1 was composed of a novel drug and a known drug, while Drugbank-S2 contained two unknown drugs.

For the metabolic DDI classification task, the MMDDI dataset comprising 343,036 DDIs was introduced^15^. Among them, each DDI describes the process by which one drug (named perpetrator) affects the PK (i.e., plasma concentrations) of the other (named victim) by inducing or inhibiting the relevant metabolic enzymes. More specifically, this dataset contained 131,376 and 40,142 drug pairs labeled as *Label*_1_ and *Label*_2_ (correct labels), respectively. After that, drug pairs with reversed semantic orders were subsequently generated, resulting in *Label*_3_ and *Label*_4_ (incorrect labels) annotations. Consequently, this task is formulated as *< D*_*a*_, *D*_*b*_ *>*→ *Label*_*i* ∈_{_1,2,3,4_}, in which the specific meaning of *Label*_*i*_ is detailed in Supplementary Table 5. The dataset possessed 1,409 unique drugs. On this basis, model performance was evaluated through five-fold cross-validation experiments. In addition, we also adopted an external test set, DDInter^45^, whose labels were composed only of *Label*_1_ and *Label*_2_, involving 2,354 and 1,012 drug pairs, respectively.

To further quantify the impact of the perpetrator drug on the PK of the victim drug, the fourth type of DDI task is introduced, i.e., PK fold change prediction. In this aspect, the AUC_FC dataset^46^ was used to evaluate model performance. This dataset was composed of 3,987 drug pairs along with the corresponding area under the plasma time-concentration curve fold change (AUC FC) values for the victim drug. To optimize the label distribution and ensure training convergence, the preprocessing operation, *log*_2_(AUC FC), was implemented. Therefore, this task can be formulated as *< D*_*a*_, *D*_*b*_ *>*→ *log*_2_(AUC FC). Similarly, model performance on this dataset was evaluated using five-fold cross-validation experiments. Furthermore, an external test set named AUC_FC_External was used to evaluate the models generalization ability. The dataset included nine drugs newly approved by the Food and Drug Administration (FDA) in 2023^47^, involving 47 DDIs.

### Node-line dual-topology drug representation

Typically, drug SMILES strings are converted into molecular graph representations using RDKit^48^, with atoms as nodes and covalent bonds as edges. Although this representation is highly consistent with the real-world molecular structure paradigm, one limitation lies in its ability to only define the first-order connectivity between atoms, making it challenging to capture global topological patterns of covalent bonds (e.g., adjacency relationships between bonds in ring-shaped functional groups). On this basis, the role of bond-level semantics in molecular representation is weakened, which contradicts the principle that both bonds and atoms are equally important for functional groups. Therefore, we proposed the concept of node-line dual-topology drug representation to fully exploit the potential of drug graph representation. As its name implies, the drug representation in this work is composed of two parts, i.e., node graph topology and line graph topology. Among them, the former refers to conventional drug graphs with atoms as nodes and bonds as edges, denoted as *G*_*n*_ = (*V*_*n*_, *E*_*n*_), where *V*_*n*_ and *E*_*n*_ represent the sets of atoms/nodes and bonds/edges, respectively. On this basis, *v*_*i*_ ∈ *V*_*n*_ means the *i*-th atom and *e*_*i, j*_ ∈ *E*_*n*_ means the bond between *i*-th and *j*-th atoms. Each atom/node *v*_*i*_ can be assigned a feature vector to define various physicochemical properties, such as atom types, valences, and others (detailed in Supplementary Table 6), resulting in the atomic feature matrix 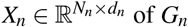. Here, *N*_*n*_ = *V*_*n*_ indicates the number of atoms and *d*_*n*_ is the dimensionality of atom features. In this way, the feature vector of *i*-th atom can be represented as *x*_*i*_ ⊂ *X*_*n*_. Furthermore, the adjacency matrix 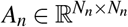 representing the connectivity among atom nodes in *G*_*n*_ can also be obtained according to the edge set *E*_*n*_.

Similarly, the line graph topology is denoted as *G*_*e*_ = (*V*_*e*_, *E*_*e*_), which can be obtained by performing the operation of topology inversion *T* (·) on the node graph, i.e., *V*_*e*_ = *T* (*E*_*n*_) and *E*_*e*_ = *T* (*V*_*n*_), as shown in Figure 1a. Specifically, bonds/edges in the node graph are mapped to nodes in the line graph, i.e., *e*_*i, j*_ ∈ *V*_*e*_., whereas atoms/nodes in the node graph are considered as edges in the line graph, i.e., *v*_*i*_ ∈ *E*_*e*_. In this way, bond-centric novel topology can be acquired to represent the structure of drug molecules. Each bond/node *e*_*i, j*_ in the line graph *G*_*e*_ can be described by a feature vector composed of bone types, conjugative statuses, and others (see Supplementary Table 7), resulting in the bond feature matrix 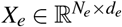. Among them, *N*_*e*_ = |*V*_*e*_| = |*E*_*n*_| is the number of bonds and *d*_*e*_ = 6 is the dimensionality of bond features. On this basis, the feature vector of *e*_*i, j*_ can be denoted as *x*_*i, j*_ ⊂ *X*_*e*_. The adjacency matrix 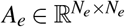 representing the connectivity among bond nodes in *G*_*e*_ can also be constructed according to edge set *E*_*e*_.

### Weave message passing neural network

After the aforementioned steps, the dual-topology drug representations composed of atom-centric node graphs and bond-centric line graphs can be constructed. To obtain a drug descriptor with powerful representational capacities, it is necessary to fuse the atom-bond semantic knowledge embedded within both topologies. Conventional operations generally employ two separate graph neural networks to extract features from each topology, followed by straightforward feature concatenation. However, this strategy fails to solve the semantic gap caused by the structural heterogeneity between the two topologies. Here, we designed the Weave Message Passing Neural Network (Weave-MPNN), as shown in Figure 9a. It alternated the feature updating processes across the node graph and line graph topologies, thereby forcing the two topologies to progressively align semantically during the iteration. Each Weave-MPNN layer logically consists of two steps, i.e., the bond-to-atom update in the node graph topology and the atom-to-bond update in the line graph topology.

**Figure 9.**
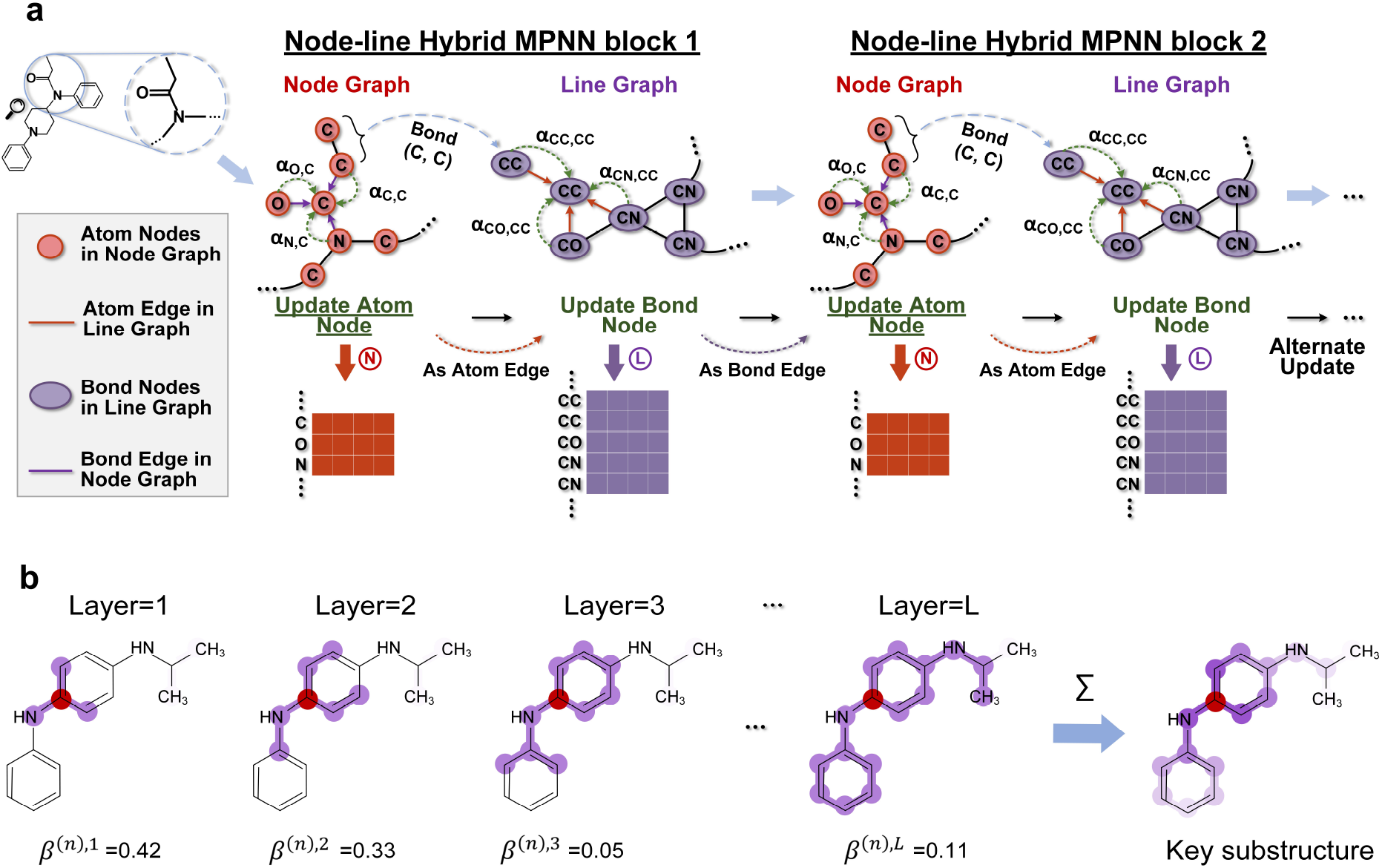
Schematic representation of Weave MPNN. **a**. The architecture of Weave MPNN. **b**. The definition of key substructures.

Specifically, each atom/node feature *x*_*i*_ in node graph *G*_*n*_ and each bond/node feature *x*_*i, j*_ in line graph *G*_*e*_ were first mapped to feature spaces with the same dimensionality through a linear layer, resulting in 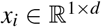 and 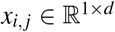, respectively. Among them, we set *d* to 64. On this basis, the bond/node feature *x*_*i, j*_ in the line graph was used to update atom/node feature *x*_*i*_ in the node graph:

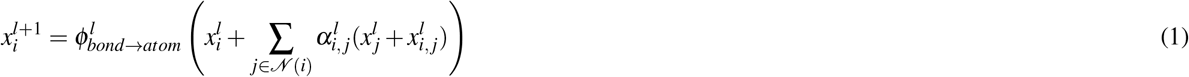

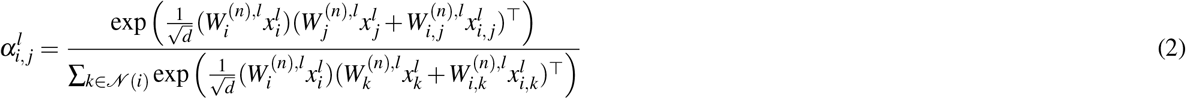

where *N* (*i*) denotes the neighbors of atom/node *v*_*i*_ in the node graph (obtained from *A*_*n*_). 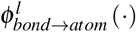 is the update function in the *l*-th layer, consisting of a linear layer and a PReLU activation function. 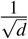 is represents the scaling factor and (·)^⊤^ the transposition. 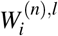 and 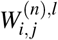 are learnable parameters specific to the atom *v*_*i*_ and bond *e*_*i j*_ in the bond-to-atom step, respectively. In this way, the attention coefficient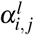 can be calculated from learnable query 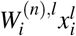and key 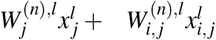 vectors. So far, the feature 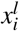 of each atom/node *v*_*i*_ in the node graph is updated to 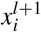.

Proceeding to the second step, we implemented atom-to-bond update in the line graph topology. In this process, the updated atom/node feature 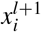 in the node graph was used to update bond/node feature 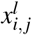 in the line graph:

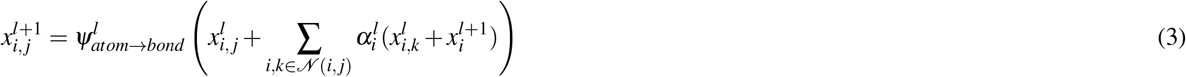

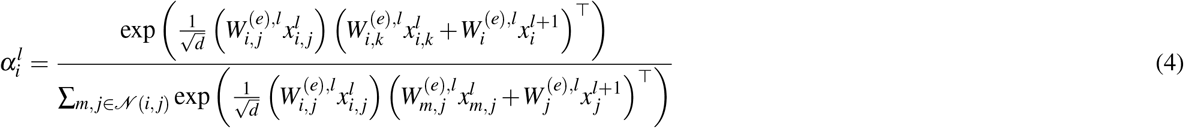

where *N* (*i, j*) denotes the neighbors of bond/node *e*_*i,j*_ in the line graph (obtained from *A*_*e*_). 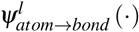 is the update function, which has the same architecture as 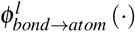, but they do not share learnable parameters. 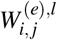 and 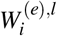 are learnable parameters specific to the bond *e*_*i,j*_ and atom *v*_*i*_ in the atom-to-bond step, respectively. Similar with the bond-to-atom step, the feature 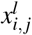 of each bond/node *e*_*i,j*_ in the line graph is updated to 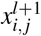. Then, 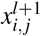 and 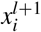 proceed to the next iteration, with the former updating the latter’s features through the bond-to-atom step. In this way, the semantic gap between the two topologies is gradually reduced during the iterations of feature updates. On this basis, 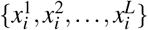 and 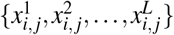 can be obtained, where *L* denotes the number of Weave-MPNN layers.

### Cross-topology consensus enhancer

As the substructure of drug molecules, functional groups are of great importance for the interaction between a pair of drugs. For example, paroxetine relies on its 1,3-benzodioxole to inhibit the activity of CYP2D6, thereby affecting the plasma concentration of another drug^27^. A frequently-used substructure extraction method involves performing convolution operations on drug graphs. The *l*-th layer convolution aggregates *l*-order neighbors, which can be considered as the extraction of drug substructures with the radius of *l*. Following this idea, atom features 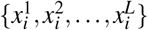 in Weave MPNN can be interpreted as 1-radius to *L*-radius substructures centered on the atom/node *v*_*i*_ in the node graph, with bond features 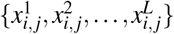 in the line graph interpreted in a similar manner. On the basis that the semantic gaps between the node-line topologies are reduced via Weave MPNN, we proposed the cross-topology consensus enhancer to reinforce the substructure consensus between atom-centric node graphs and bond-centric line graphs, enabling the synergistic localization of key substructures within drug molecules.

First, we tried to assign different attention coefficients β to substructures with different radii in the node graph and line graph. To achieve this, the graph-level representations of the two graph topologies in the *l*-th layers, *g*^(*n*),*l*^ and *g*^(*e*),*l*^,were separately calculated using the SAGPooling with a pooling ratio of 1:

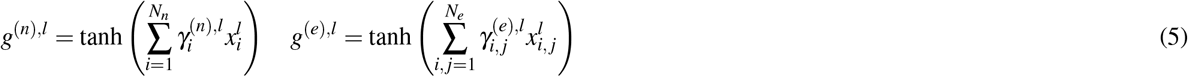

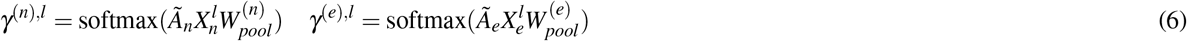

where 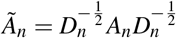 and 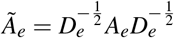 are symmetrically normalized adjacency matrixes of the node graph and line graph. Similarly, 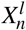 and 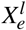 are feature matrixes in the *l*-th layer of Weave MPNN, which is the vertical concatenation of all atom feature 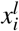 and all bond feature 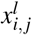. 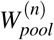and 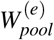 are learnable parameters used to calculate pooling attention matrixes 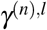 and 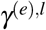 in the node graph and line graph, respectively. Therefore, 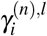 and 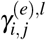 are the pooling attentions of atom *v*_*i*_ and bond *e*_*i,j*_ in the *l*-th layer. Afterwards, taking the node graph as an example, we calculated the attention coefficients β^(*n*)^ assigned to 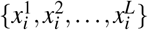 through different layers of graph-level representations *g*^(*n*)^, which can be defined as follows:

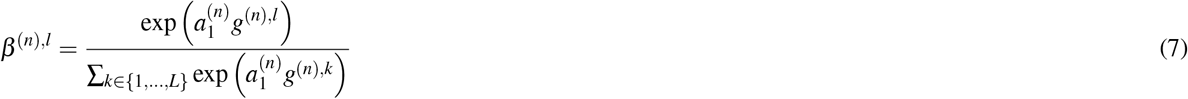

Here, 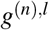 denotes the graph-level representation in the *l*-th layer, and *L* denotes the number of Weave-MPNN layers. 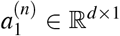 indicates the learnable projection vector, which is used to transform *g*^(*n*),*l*^ into a real number. On this basis, the substructures with different radii were summed according to their attention coefficients, resulting in the key substructure *s*_*i*_ (see Figure 9b) centered on atom *v*_*i*_:

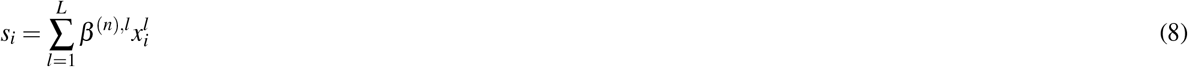

Following the same steps as the node graph, the attention coefficients β^(*e*)^ assigned to 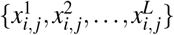 and the key substructures *s*_*i, j*_ centered on bond *e*_*i, j*_ in the line graph can be obtained. It is noteworthy that β^(*n*)^ and β^(*e*)^ are crucial for determining the key substructures in the node graph and line graph, respectively. However, they are independent in the above calculation process, which may lead to different substructure insights in the two topologies of a drug. To this end, we imposed consistency constraints on the graph-level representations *g*^(*n*)^ and *g*^(*e*)^ to enhance substructure consensus between the two topologies:

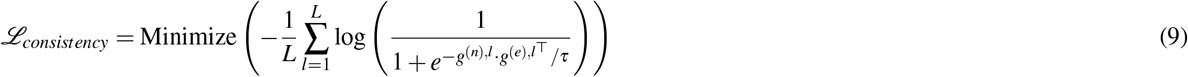

where τ is a temperature parameter set to 0.2. Finally, through a projection function *f* (·), the key substructure *s*_*i*_ extracted by atom *v*_*i*_ was projected back into its atom/node representation *m*_*i*_ in the node graph, thereby injecting substructural information into *m*_*i*_:

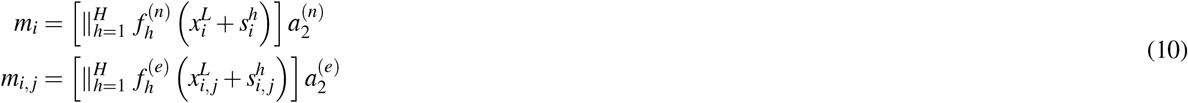

Here, we applied multi-head mechanism. *H* denotes the number of heads, and *f* (·) implements the architecture of multi-layer perceptron. 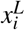 is the maximum-radius substructure that can be extracted by *v*_*i*_, which supplements global information for *m*_*i*_. *a*_2_ ∈ R^*Hd*×*d*^ is the learnable projection vector used to recover the original dimensionality *d*. Similarly, bond/node representation *m*_*i, j*_ in the line graph can be obtained.

### Cross-entity interaction module

Among a pair of drugs intend for interaction prediction, the first drug is able to acquire its atom/node representations 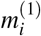 in the node graph and bond/node representations 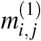 in the line graph according to the above steps. Identically, 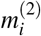 and 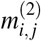 for another drug are availability as well. In this section, we proposed the cross-entity interaction module, which calculated atom-wise affinities and bond-wise affinities (i.e., attractive forces) between a pair of drugs from the perspectives of the node graph and line graph, respectively. This enabled systematic simulation of the interaction process between two drug entities. On this basis, atoms or bonds with strong affinities to another drug can be identified, thereby endowing the model with dual-topology molecular interpretability. Specifically, atom-wise affinities and bond-wise affinities between a pair of drugs can be separately defined as:

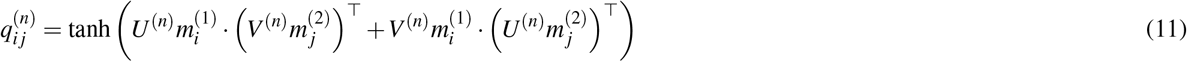

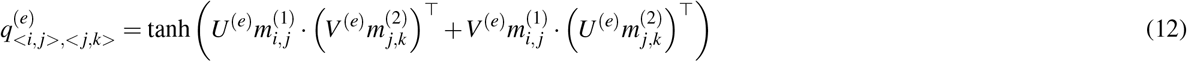

where 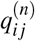 denotes the affinity between *i*-th atom in the first drug and *j*-th atom in the second drug. Similarly, 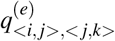 indicates the affinity between bond *e*_*i, j*_ in the first drug and bond *e* _*j,k*_ in the second drug. *U* ∈ R^*d*×*d*^ and *V* ∈ R^*d*×*d*^ are learnable linear transformation. After that, we can calculate the overall affinity of each atom or bond in a drug to another drug by performing the column-wise and row-wise averaging operation on 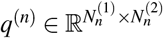 and 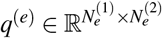:

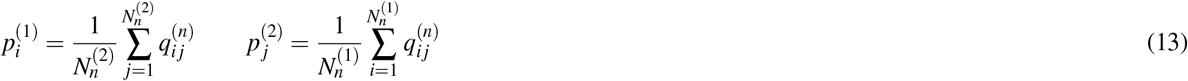

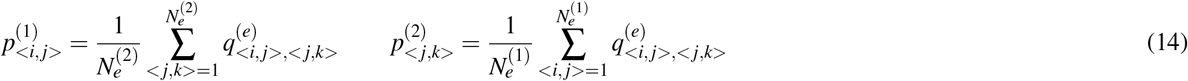

where 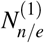 and 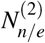 are the number of atoms/bonds in the pair of drugs, respectively. 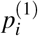 means the overall affinity of *i*-th and atom in the first drug to the second drug, and 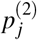 means that of *j*-th atom in the second drug to the first drug. Similarly, 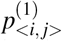 and 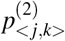 can be explained as well. Therefore, the atoms or bonds with greater affinity values to another drug can be observed. Furthermore, these affinities were normalized to the range from 0 to 1 via the softmax function to quantify the importance of the respective atoms or bonds in drug interactions:

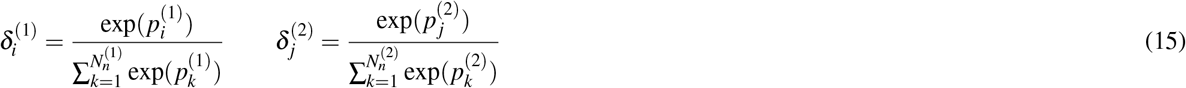

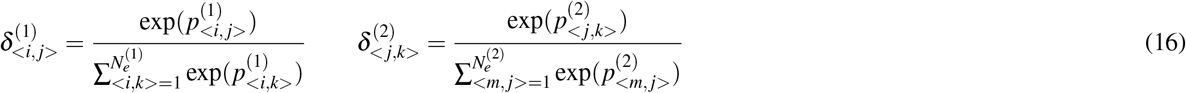

In this way, by visualizing the importance scores 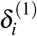 and 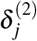 in the node graph, along with 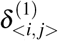 and 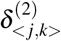 in the line graph, critical atoms and bonds between the pair of drugs can be precisely localized. In other words, node-line dual-topology representations jointly elucidated the molecular interpretability of the model. On this basis, graph-level representations of node graph and line graph in the pair of drugs, *z*_*n*_ ∈ R^1×*d*^ and *z*_*e*_ ∈ R^1×*d*^, can be obtained:

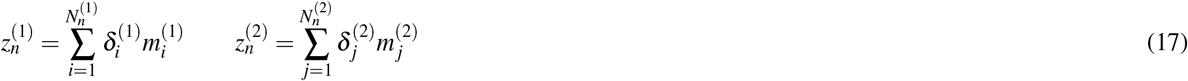

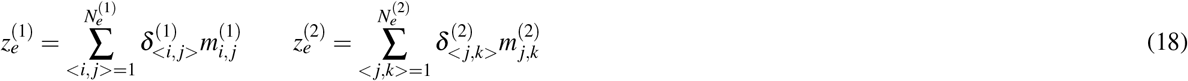

Finally, dual-topology graph-level representations for the pair of drugs were horizontally concatenated and subsequently fed into four different multilayer perceptron decoder *f*_1∼4_(·) for the DDI prediction, each corresponding to one of the four DDI task types. Among them, *f*_1∼2_(·) was followed by the sigmoid function for *task*_1™2_, whereas *f*_3_(·) was followed by the softmax function for *task*_3_. Noteworthily, *task*_2_ required the DDI type to be input, so the DDI prediction for *task*_2_ can be defined as 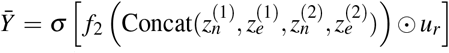, where *u*_r_ ∈ R^1×*d*^ denotes the learnable representation of DDI type *r*, and σmeans the sigmoid function.

### Experimental settings

In the proposed model, Adam^49^ is chosen as the optimizer. Due to the differences in the type of DDI tasks and the size of the datasets, the model adopts different loss functions *L*_*task* ∈_{_1,2,3,4_} and learning rates for different DDI tasks. In the first two types of DDI tasks, the loss function and learning rate are set to binary cross-entropy (BCE) and 0.001, respectively. However, the loss functions of the third and fourth types of DDI tasks are set to cross-entropy (CE) and mean square error (MSE), respectively, where their learning rates are both 0.0001. Therefore, the final loss function used for backpropagation is defined as *L* = λ_1_*L*_*task*_ + λ_2_*L*_*consistency*_, where λ_1_ and λ_2_ range from 0 to 1. In addition, the calculation of atom/bond-wise affinity 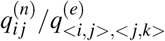 in the cross-entity interaction module applied the operation of transpose multiplication. On this basis, we tried other two operations as well, i.e., matrix addition and Hadamard product. The above three different operations are taken as the critical hyperparameters together with the number of Weave MPNN layers, the number of heads in cross-topology consensus enhancer, λ_1_ and λ_2_. After hyperparameter optimization (Figure 7c and Figure 8), we still end up using the transpose multiplication to compute atom/bond-wise affinity. Similarly, the number of Weave MPNN layers is set to 10, the number of heads is set to 2. λ_1_ and λ_2_ are set to 1 and 0.4, respectively. Furthermore, six metrics—Accuracy (Accuracy), Area Under the ROC Curve (AUC), F1-Score (F1), Precision (Prec), Recall (Rec), and Average Precision (AP)—were employed to evaluate the model’s performance on DDI classification tasks. For DDI regression tasks, five metrics—Root Mean Squared Error (RMSE), Mean Squared Error (MSE), Pearson Correlation Coefficient (Pearson), Spearman Rank Correlation Coefficient (Spearman), and Concordance Index (CI)—were used to assess the model’s performance. Finally, the proposed model and related experiments were implemented using PyTorch^50^ on an NVIDIA GeForce RTX 4090 graphics card with ∼24GB memory.

### Simulations using virtual heart

The geometry of the virtual heart was reconstructed from DT-MRI^51^. The geometry was then discretized into approximately 24 million voxels with each of them being assigned with a mathematical cell model of human ventricular myocyte. The Ohara Rudy dynamic (ORd) cell model is used in this study. To model the effect of quinidine, we incorporated all its known actions on cardiac ionic currents including *I*_Kr_, *I*_Ks_, *I*_K1_, *I*_to_, *I*_CaL_, *I*_Na_, and *I*_NaL_^52^. The concentration-dependent behaviour was modeled based on the pore block theory, and a serum concentration of 5 µmol/L was applied to reflect a therapeutic usage. When it was co-administrated with ritonavir, the concentration of quinidine was increased to 15 µmol/L to mimic the condition where its metabolism via CYP3A4 was inhibited by ritonavir. These cells with the drug effects incorporated were then coupled using a monodomain equation to describe the propagation of excitation waves in the three-dimensional heart^53^:

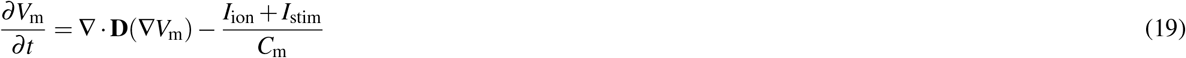

where *V*_*m*_ denotes the membrane potential, *I*_ion_ and *I*_stim_ are the total ionic current and the stimulating current, respectively, and *C*_*m*_ is the membrane capacitance. *D* is the diffusion coefficient tensor for describing the intercellular electrical coupling via gap junctions. The pseudo-ECG was generated using the following equations:

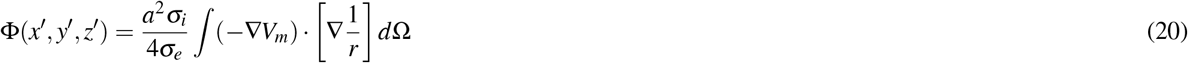

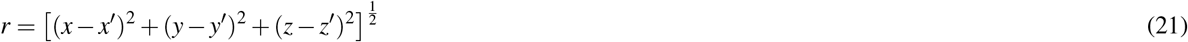

where Φ is a unipolar potential generated from the ventricular model, *r* denotes the distance from a source site on the ventricle to the virtual electrode. σ_*i*_ and σ_*e*_ are intracellular and extracellular conductivities, respectively.

### Comparison methods

For benchmarking, we implemented 9 state-of-the-art methods for DDI prediction. The details of the 9 SOTA methods are as follows:

1. DeepDDI^42^: DeepDDI computes structural similarity profiles of drug pairs and inputs them into a deep neural network to predict whether an interaction occurs between the pair.
2. GAT-DDI^54^: GAT-DDI employs multiple graph attention network (GAT) layers to dynamically learn complex atomic relationships within drug molecules, thereby extracting structural information for DDI prediction.
3. SAGE-DDI^55^: Similar to GAT-DDI, SAGE-DDI adopts the graph sample and aggregation (GraphSAGE) algorithm as its architecture backbone for DDI prediction.
4. MR-GNN^56^: MR-GNN designs dual LSTM architectures: Summary-LSTM for aggregating multi-resolution local features and Interaction-LSTM for extracting interaction features between drug pairs, thereby enhancing DDI prediction.
5. SSI-DDI^18^: SSI-DDI directly learns drug substructures from molecular graphs and quantifies substructural contributions to DDIs through a designed co-attention mechanism.
6. DGNN-DDI^57^: Compared with SSI-DDI, DGNN-DDI designs a directed message passing neural network with sub-structure attention mechanism (SA-DMPNN), which also achieves superior DDI prediction performance.
7. SA-DDI^16^: Similarly, SA-DDI extracts flexible-sized and irregular-shaped substructures and predicts drug interactions by modeling chemical reactions between these substructures.
8. SRR-DDI^17^: SRR-DDI calculates drug similarity features and substructure features, then derives final drug representations via a dual drug feature fusion module for DDI prediction.
9. MeTDDI^15^: MeTDDI represents drugs as motif-based graphs and achieves superior DDI prediction performance through a Transformer encoder with local-global self-attention and a co-attention network.

## Data availability

The datasets used for the model construction are available at https://pan.baidu.com/s/1-i8GCBnG1yFElVAg6bviiA?pwd=1234. The DDI database predicted by DualTopoDDI is available at https://github.com/Wenjian-Ma/DualTopoDDI. The Drugbank Database is available at https://go.drugbank.com/releases/latest.

## Code availability

The code is available at https://github.com/Wenjian-Ma/DualTopoDDI. The software used in this work is as follows: python (3.8.17), pytorch (1.13.1), PyG (2.2.0), sklearn (1.3.0), scipy (1.9.3), numpy (1.24.3), and RDKit (2023.9.6).

## Acknowledgements

This work was supported by the National Natural Science Foundation of China (NO.62306293), the Fundamental Research Funds for the Central Universities (NO.202461008; NO.202561013), and the 111 Project (NO. B23048).

## Author contributions statement

Wenjian Ma, Zhiqiang Wei, Henggui Zhang, and Shugang Zhang conceived the concept. Wenjian Ma, Xiangpeng Bi, Huasen Jiang, Weigang Lu, Jiaxin Lin, and Shutan Lin designed the methodology and performed the experiments. Wenjian Ma, Jie Nie, and Shugang Zhang analyzed the results. Wenjian Ma and Shugang Zhang wrote the manuscript with the help from all authors. All authors reviewed and approved the final version of this manuscript.

