## Supplementary Information for "Deciphering Mechanistic Signatures in Drug-Drug Interactions with Dual Topology Graphs"

### Contents

1. Supplementary Tables 1 to 7.

**Supplementary Table 1.** Performance comparison (mean±std in %) of DualTopoDDI with baseline methods on the standard DDI task.

| Datasets | Methods | ACC↑ | AUC↑ | F1↑ | Prec↑ | Rec↑ | AP↑ |
| --- | --- | --- | --- | --- | --- | --- | --- |
| DeepDDI | DeepDDI | 90.99±0.85 | 94.76±1.30 | 93.82±0.66 | 97.30±0.53 | 90.60±1.66 | 97.31±0.66 |
|  | GAT-DDI | 87.01±0.33 | 90.64±0.03 | 91.40±0.26 | 92.11±0.25 | 90.72±0.76 | 95.99±0.04 |
|  | SAGE-DDI | 88.86±0.19 | 92.62±0.28 | 92.64±0.13 | 93.21±0.24 | 92.08±0.29 | 96.85±0.08 |
|  | MR-GNN | 90.93±0.01 | 99.53±0.01 | 93.63±0.01 | <u>99.62±0.01</u> | 88.23±0.01 | 99.83±0.01 |
|  | SSI-DDI | 92.72±0.27 | 97.53±0.03 | 95.11±0.18 | 97.33±0.09 | 92.99±0.40 | 99.06±0.01 |
|  | DGNN-DDI | 97.88±0.01 | 99.64±0.01 | 98.60±0.01 | 99.30±0.03 | 97.90±0.02 | 99.87±0.01 |
|  | SA-DDI | 98.21±0.09 | 99.62±0.03 | 98.81±0.06 | 99.28±0.05 | 98.36±0.08 | 99.86±0.01 |
|  | SRR-DDI | 98.47±0.03 | <u>99.78±0.01</u> | 98.98±0.02 | 99.39±0.03 | <u>98.58±0.03</u> | 99.92±0.01 |
|  | MeTDDI | <u>98.49±0.10</u> | 99.78±0.03 | <u>99.00±0.07</u> | 99.56±0.05 | 98.44±0.10 | <u>99.92±0.01</u> |
|  | **Ours** | **98.98±0.04** | **99.84±0.01** | **99.33±0.03** | **99.65±0.01** | **99.01±0.05** | **99.93±0.01** |
| ZhangDDI | DeepDDI | 87.68±0.90 | 90.46±0.29 | 74.37±1.90 | 70.40±1.72 | 78.82±2.18 | 71.32±2.41 |
|  | GAT-DDI | 88.32±0.06 | 93.39±0.06 | 74.66±0.36 | 73.52±1.00 | 75.88±1.83 | 80.75±0.40 |
|  | SAGE-DDI | 89.74±0.44 | 94.71±0.21 | 77.85±1.03 | 76.29±0.84 | 79.47±1.34 | 84.69±0.63 |
|  | MR-GNN | 81.51±0.21 | 96.03±0.05 | 70.39±0.29 | 55.29±0.27 | **96.84±0.26** | 84.74±0.67 |
|  | SSI-DDI | 92.67±0.12 | 97.17±0.08 | 83.93±0.26 | 83.52±1.64 | 84.42±2.01 | 92.70±0.10 |
|  | DGNN-DDI | 94.06±0.19 | 98.01±0.17 | 86.91±0.49 | <u>86.91±0.33</u> | 86.92±1.01 | 94.90±0.38 |
|  | SA-DDI | 93.44±0.22 | 97.66±0.07 | 85.69±0.62 | 84.84±0.22 | 86.59±1.50 | 94.18±0.30 |
|  | SRR-DDI | <u>94.09±0.19</u> | <u>98.12±0.13</u> | <u>87.06±0.35</u> | 86.59±2.19 | 87.65±2.35 | <u>95.23±0.26</u> |
|  | MeTDDI | 93.25±0.10 | 97.90±0.06 | 85.69±0.16 | 82.78±0.50 | 88.84±0.26 | 94.62±0.17 |
|  | **Ours** | **94.73±0.28** | **98.48±0.11** | **88.45±0.60** | **88.03±0.72** | <u>88.87±0.51</u> | **96.11±0.33** |
| ChChMiner | DeepDDI | 91.17±0.49 | 96.48±0.34 | 94.51±0.66 | 98.18±1.61 | 91.13±0.29 | 99.02±0.52 |
|  | GAT-DDI | 91.32±0.37 | 96.51±1.12 | 94.94±0.21 | 98.06±1.09 | <u>92.03±0.89</u> | 99.51±0.16 |
|  | SAGE-DDI | 90.78±0.17 | 96.91±0.99 | 94.58±0.14 | 98.52±0.69 | 90.96±0.85 | 99.58±0.13 |
|  | MR-GNN | 78.44±0.01 | 98.54±0.01 | 86.13±0.01 | 99.59±0.01 | 75.64±0.01 | 99.74±0.01 |
|  | SSI-DDI | 91.84±0.39 | 98.64±0.19 | 95.19±0.24 | 99.55±0.01 | 91.19±0.43 | 99.82±0.02 |
|  | DGNN-DDI | 90.82±0.26 | <u>99.36±0.11</u> | 94.54±0.16 | <u>99.78±0.08</u> | 89.83±0.31 | <u>99.91±0.01</u> |
|  | SA-DDI | 90.65±0.47 | 99.06±0.08 | 94.43±0.29 | 99.75±0.08 | 89.65±0.58 | 99.85±0.04 |
|  | SRR-DDI | 91.11±0.39 | 99.11±0.06 | 94.72±0.24 | 99.65±0.03 | 90.25±0.41 | 99.88±0.01 |
|  | MeTDDI | **96.01±0.27** | **99.79±0.06** | **97.69±0.16** | **99.97±0.02** | **95.51±0.30** | **99.97±0.01** |
|  | **Ours** | <u>92.27±0.30</u> | 99.22±0.14 | <u>95.45±0.18</u> | 99.66±0.07 | 91.57±0.40 | 99.89±0.02 |

Bold and underline indicate the optimal and suboptimal performance, respectively.

**Supplementary Table 2.** Performance comparison (mean±std in %) of DualTopoDDI with baseline methods on the DDI type prediction task.

| Datasets | Methods | ACC↑ | AUC↑ | F1↑ | Prec↑ | Rec↑ | AP↑ |
| --- | --- | --- | --- | --- | --- | --- | --- |
| DrugBank | DeepDDI | 93.21±0.27 | 97.03±0.11 | 93.37±0.22 | 91.26±0.26 | 95.52±0.43 | 95.95±0.21 |
|  | GAT-DDI | 92.15±0.11 | 96.19±0.12 | 92.29±0.16 | 90.28±0.19 | 95.34±0.33 | 95.02±0.03 |
|  | SAGE-DDI | 83.05±0.08 | 89.96±0.09 | 83.73±0.21 | 80.52±0.72 | 87.23±1.26 | 87.22±0.13 |
|  | MR-GNN | 93.26±0.14 | 97.26±0.04 | 93.35±0.12 | 91.26±0.21 | 95.69±0.02 | 96.45±0.07 |
|  | SSI-DDI | 94.24±0.13 | 98.12±0.03 | 94.29±0.08 | 93.01±0.02 | 96.11±0.07 | 97.25±0.02 |
|  | DGNN-DDI | 96.12±0.05 | 98.90±0.02 | 95.98±0.16 | 94.86±0.07 | 97.89±0.03 | 98.46±0.05 |
|  | SA-DDI | 96.21±0.13 | 98.78±0.04 | 96.27±0.11 | 95.01±0.08 | 97.62±0.02 | 98.36±0.03 |
|  | SRR-DDI | <u>96.67±0.06</u> | <u>99.05±0.03</u> | <u>96.72±0.05</u> | <u>95.28±0.08</u> | <u>98.24±0.03</u> | <u>98.74±0.04</u> |
|  | MeTDDI | 96.19±0.06 | 98.91±0.02 | 96.21±0.04 | 94.90±0.09 | 97.57±0.02 | 98.61±0.05 |
|  | **Ours** | **97.51±0.08** | **99.39±0.04** | **97.54±0.09** | **96.28±0.09** | **98.82±0.08** | **99.08±0.05** |
| TWO SIDES | DeepDDI | 75.16±0.23 | 82.42±0.31 | 77.03±0.05 | 71.65±0.59 | 83.27±0.84 | 79.47±0.33 |
|  | GAT-DDI | 67.32±2.04 | 75.16±2.44 | 63.70±3.11 | 71.54±2.19 | 57.65±5.09 | 72.48±2.45 |
|  | SAGE-DDI | 81.02±0.08 | 88.15±0.04 | 82.22±0.08 | 77.35±0.12 | 87.75±0.19 | 84.85±0.09 |
|  | MR-GNN | 85.39±0.31 | 91.93±0.21 | 86.46±0.27 | 80.57±0.37 | 93.28±0.21 | 89.32±0.22 |
|  | SSI-DDI | 82.21±0.38 | 89.25±0.45 | 83.32±0.45 | 79.15±0.32 | 87.55±0.64 | 86.19±0.41 |
|  | DGNN-DDI | 85.33±0.12 | 92.09±0.12 | 84.87±0.31 | 81.03±0.17 | 89.35±0.17 | 89.78±0.24 |
|  | SA-DDI | 87.45±0.06 | 93.15±0.04 | <u>88.35±0.04</u> | 82.43±0.02 | <u>95.18±0.10</u> | <u>90.51±0.08</u> |
|  | SRR-DDI | <u>87.52±0.05</u> | <u>93.20±0.07</u> | 88.32±0.05 | **83.06±0.02** | 94.38±0.11 | 90.37±0.06 |
|  | MeTDDI | 85.42±0.09 | 92.05±0.13 | 86.38±0.11 | 81.07±0.03 | 92.43±0.05 | 89.58±0.09 |
|  | **Ours** | **87.94±0.02** | **93.51±0.02** | **88.82±0.02** | <u>82.74±0.03</u> | **95.87±0.02** | **91.04±0.02** |
| Drugbank-S1 | DeepDDI | 74.80±0.38 | 79.48±0.85 | 73.08±1.13 | 78.44±1.28 | 68.52±2.87 | 77.16±0.87 |
|  | GAT-DDI | 73.00±0.54 | 79.99±0.39 | 71.74±0.82 | 75.23±0.81 | 68.60±1.72 | 78.67±0.37 |
|  | SAGE-DDI | 73.64±0.22 | 80.37±0.25 | 72.64±0.45 | 75.55±1.31 | <u>70.01±1.92</u> | 79.38±0.49 |
|  | MR-GNN | <u>76.98±0.26</u> | <u>84.32±0.41</u> | <u>75.58±0.25</u> | 80.50±0.46 | **71.22±0.34** | <u>83.75±0.25</u> |
|  | SSI-DDI | 74.70±0.22 | 82.02±0.41 | 72.95±0.42 | 78.37±0.21 | 68.23±0.85 | 81.89±0.42 |
|  | DGNN-DDI | 74.47±0.11 | 82.11±0.36 | 71.92±1.50 | 80.20±3.70 | 65.65±5.26 | 82.57±1.04 |
|  | SA-DDI | 75.54±0.47 | 83.55±0.15 | 72.60±1.53 | 82.61±2.44 | 65.00±4.04 | 83.66±0.63 |
|  | SRR-DDI | 75.76±0.41 | 83.44±0.25 | 73.90±0.86 | 80.04±1.07 | 68.69±2.14 | 82.88±0.19 |
|  | MeTDDI | 73.61±0.25 | 80.78±0.59 | 69.12±0.64 | <u>83.35±1.65</u> | 59.09±1.66 | 82.13±0.70 |
|  | **Ours** | **78.41±0.21** | **85.68±0.18** | **75.95±0.34** | **85.76±1.41** | 68.19±1.35 | **86.89±0.24** |
| Drugbank-S2 | DeepDDI | 65.65±0.79 | 69.46±1.63 | 57.37±2.60 | 75.57±1.30 | 46.40±3.76 | 69.55±0.73 |
|  | GAT-DDI | 66.58±0.48 | 73.10±0.21 | <u>60.99±1.43</u> | 73.25±1.05 | 42.32±2.53 | 72.44±0.10 |
|  | SAGE-DDI | 66.75±0.71 | 72.76±0.28 | **61.66±1.76** | 72.81±1.33 | 43.59±3.16 | 72.71±0.48 |
|  | MR-GNN | 67.06±0.33 | 73.46±0.39 | 58.55±0.34 | 71.12±0.47 | 47.44±0.29 | 73.54±0.28 |
|  | SSI-DDI | 66.18±0.55 | 73.17±0.87 | 58.72±1.12 | 75.33±0.62 | <u>48.14±1.52</u> | 74.07±0.89 |
|  | DGNN-DDI | 65.43±2.01 | 72.65±1.90 | 55.27±6.70 | 78.16±3.80 | 43.92±6.13 | 74.55±0.88 |
|  | SA-DDI | 66.49±1.36 | 74.80±0.35 | 57.65±4.29 | <u>78.22±2.39</u> | 46.08±5.90 | 75.53±0.04 |
|  | SRR-DDI | <u>67.21±0.83</u> | <u>75.66±0.29</u> | 59.83±2.23 | 77.18±1.07 | **48.97±3.41** | <u>76.16±0.61</u> |
|  | MeTDDI | 63.11±0.05 | 68.95±1.18 | 55.58±3.25 | 70.27±3.69 | 46.62±6.31 | 69.63±1.39 |
|  | **Ours** | **67.56±0.31** | **75.70±0.07** | 58.76±1.60 | **80.68±1.94** | 46.32±2.68 | **77.59±0.32** |

Bold and underline indicate the optimal and suboptimal performance, respectively.

**Supplementary Table 3.** Performance comparison (mean±std in %) of DualTopoDDI with baseline methods on the metabolic DDI classification task.

| Datasets | Methods | ACC↑ | AUC↑ | F1↑ | Prec↑ | Rec↑ | AP↑ |
| --- | --- | --- | --- | --- | --- | --- | --- |
| MMDDI | DeepDDI | 78.91±0.27 | 94.87±0.06 | 78.50±0.33 | 78.04±0.51 | 79.12±0.49 | 85.92±0.40 |
|  | GAT-DDI | 77.36±0.27 | 94.39±0.11 | 76.48±0.35 | 77.22±0.41 | 75.88±0.88 | 84.85±0.31 |
|  | SAGE-DDI | 79.81±0.07 | 95.41±0.03 | 79.13±0.12 | 79.41±0.44 | 78.92±0.62 | 87.56±0.10 |
|  | MR-GNN | 87.96±0.24 | 98.08±0.05 | 88.22±0.25 | 88.48±0.17 | 87.97±0.34 | 94.87±0.15 |
|  | SSI-DDI | 81.92±0.41 | 96.34±0.15 | 81.80±0.59 | 82.03±0.68 | 81.61±0.60 | 90.23±0.54 |
|  | DGNN-DDI | <u>95.27±0.38</u> | <u>99.63±0.03</u> | <u>94.87±0.46</u> | <u>94.86±0.49</u> | <u>94.88±0.42</u> | <u>98.83±0.14</u> |
|  | SA-DDI | 92.56±0.28 | 99.29±0.04 | 92.57±0.34 | 92.49±0.34 | 92.65±0.36 | 98.01±0.14 |
|  | SRR-DDI | 93.42±0.18 | 99.43±0.02 | 93.47±0.21 | 93.50±0.21 | 93.44±0.22 | 98.41±0.08 |
|  | MeTDDI | 93.21±0.01 | 99.36±0.01 | 93.25±0.04 | 93.32±0.05 | 93.18±0.05 | 98.18±0.03 |
|  | **Ours** | **97.90±0.06** | **99.83±0.01** | **97.86±0.09** | **97.87±0.10** | **97.84±0.09** | **99.45±0.03** |
| DDInter | DeepDDI | 61.64±1.38 | 83.39±0.93 | 34.45±0.68 | 40.46±1.11 | <u>30.07±0.99</u> | 39.98±1.19 |
|  | GAT-DDI | 62.02±0.63 | 84.17±0.44 | 34.52±0.60 | 42.38±0.57 | 29.35±0.68 | 41.52±0.35 |
|  | SAGE-DDI | 61.18±1.20 | 83.90±0.70 | 34.13±0.93 | 41.67±0.59 | 29.09±1.02 | 41.37±0.68 |
|  | MR-GNN | 48.14±0.33 | 75.13±0.84 | 30.14±0.23 | <u>43.11±0.34</u> | 23.23±0.22 | 40.68±0.39 |
|  | SSI-DDI | 62.07±0.96 | <u>84.89±0.88</u> | <u>35.45±0.75</u> | 42.68±0.59 | 29.40±0.92 | <u>42.95±0.87</u> |
|  | DGNN-DDI | 61.23±0.89 | 84.88±0.68 | 34.38±0.50 | 42.17±0.61 | 29.20±0.64 | 42.33±0.62 |
|  | SA-DDI | <u>62.15±0.55</u> | 84.85±0.75 | 34.70±0.58 | 42.67±0.25 | 29.49±0.64 | 42.08±0.54 |
|  | SRR-DDI | 61.70±0.68 | 84.66±0.46 | 34.71±0.53 | 41.87±0.60 | 29.82±0.95 | 41.56±0.54 |
|  | MeTDDI | 60.30±0.35 | 83.52±0.18 | 33.73±0.31 | 42.00±0.34 | 28.42±0.30 | 41.12±0.24 |
|  | **Ours** | **62.46±0.52** | **85.09±0.72** | **35.69±0.24** | **44.02±1.08** | **30.21±0.44** | **43.24±0.53** |

Bold and underline indicate the optimal and suboptimal performance, respectively.

**Supplementary Table 4.** Performance comparison (mean±std in %) of DualTopoDDI with baseline methods on the PK fold change prediction task.

| Datasets | Methods | RMSE↓ | MSE↓ | Pearson↑ | Spearman↑ | CI↑ |
| --- | --- | --- | --- | --- | --- | --- |
| AUC_FC | DeepDDI | 0.784±0.054 | 0.617±0.087 | 0.540±0.032 | 0.407±0.046 | 0.666±0.020 |
|  | GAT-DDI | 0.723±0.037 | 0.524±0.054 | 0.561±0.043 | 0.452±0.049 | 0.684±0.021 |
|  | SAGE-DDI | 0.724±0.046 | 0.526±0.068 | 0.559±0.035 | 0.445±0.039 | 0.681±0.015 |
|  | MR-GNN | 0.764±0.045 | 0.587±0.070 | 0.582±0.031 | 0.482±0.020 | 0.698±0.009 |
|  | SSI-DDI | 0.680±0.028 | 0.463±0.038 | 0.632±0.036 | 0.532±0.037 | 0.720±0.017 |
|  | DGNN-DDI | 0.633±0.024 | 0.401±0.031 | 0.694±0.031 | 0.545±0.029 | 0.728±0.013 |
|  | SA-DDI | 0.631±0.025 | 0.399±0.032 | 0.695±0.026 | 0.542±0.034 | 0.726±0.015 |
|  | SRR-DDI | <u>0.618±0.010</u> | <u>0.382±0.012</u> | <u>0.712±0.044</u> | <u>0.566±0.036</u> | <u>0.737±0.015</u> |
|  | MeTDDI | 0.630±0.031 | 0.398±0.040 | 0.693±0.023 | 0.561±0.035 | 0.706±0.014 |
|  | **Ours** | **0.604±0.023** | **0.365±0.027** | **0.731±0.025** | **0.598±0.028** | **0.753±0.012** |
| AUC_FC<br>_External | DeepDDI | 1.048±0.076 | 1.106±0.162 | 0.671±0.063 | 0.586±0.040 | 0.716±0.019 |
|  | GAT-DDI | 1.112±0.029 | 1.238±0.064 | 0.565±0.032 | 0.532±0.056 | 0.693±0.023 |
|  | SAGE-DDI | 1.106±0.023 | 1.225±0.051 | 0.584±0.014 | 0.566±0.027 | 0.702±0.012 |
|  | MR-GNN | 1.070±0.077 | 1.152±0.161 | 0.640±0.071 | 0.595±0.092 | 0.725±0.042 |
|  | SSI-DDI | 0.972±0.067 | 0.949±0.130 | 0.720±0.051 | 0.706±0.045 | 0.769±0.023 |
|  | DGNN-DDI | 0.849±0.119 | 0.735±0.186 | 0.746±0.026 | 0.698±0.069 | 0.776±0.019 |
|  | SA-DDI | 0.908±0.038 | 0.827±0.071 | 0.757±0.018 | 0.706±0.037 | 0.770±0.020 |
|  | SRR-DDI | 0.866±0.082 | 0.757±0.145 | 0.773±0.046 | 0.702±0.056 | 0.771±0.028 |
|  | MeTDDI | <u>0.826±0.056</u> | <u>0.686±0.097</u> | <u>0.802±0.021</u> | <u>0.770±0.010</u> | <u>0.786±0.024</u> |
|  | **Ours** | **0.727±0.087** | **0.536±0.129** | **0.844±0.049** | **0.797±0.041** | **0.820±0.023** |

Bold and underline indicate the optimal and suboptimal performance, respectively.

**Supplementary Table 5.** The specific meaning of labels in the metabolic DDI classification task

| Labels | Meanings |
| --- | --- |
| *Label*<sub>1</sub> | The metabolism of Drug 1 can be decreased when combined with Drug 2. |
| *Label*<sub>2</sub> | The metabolism of Drug 1 can be increased when combined with Drug 2. |
| *Label*<sub>3</sub> | The metabolism of Drug 2 can be decreased when combined with Drug 1. |
| *Label*<sub>4</sub> | The metabolism of Drug 2 can be increased when combined with Drug 1. |

**Supplementary Table 6.** The statistic of atom features

| Atom features | Descriptions | Dims |
| --- | --- | --- |
| Atom symbols | Atom types | Total number of atom types in the dataset |
| Hybridization | [sp, sp2, sp3, sp3d, sp3d2] | 5 |
| Degree | The number of adjacent atoms | 11 |
| Implicit valence | [0, 1, 2, 3, 4, 5, 6,] | 7 |
| Formal charge | Formal charge of the atom | 1 |
| Aromatic | The atom is aromatic or not | 1 |
| Radical electrons | The number of radical electrons for the atom | 1 |

**Supplementary Table 7.** The statistic of bond features

| Bond features | Descriptions | Dims |
| --- | --- | --- |
| Bond type | [single, double, triple, aromatic] | 4 |
| Conjugated | The bond is part of a conjugated system or not | 1 |
| Ring | The bond is part of a ring or not | 1 |
